# CRISPR-Cas12a exhibits metal-dependent specificity switching

**DOI:** 10.1101/2023.11.29.569287

**Authors:** Giang T. Nguyen, Michael A. Schelling, Kathryn A. Buscher, Aneisha Sritharan, Dipali G. Sashital

## Abstract

Cas12a is the immune effector of type V-A CRISPR-Cas systems and has been co-opted for genome editing and other biotechnology tools. The specificity of Cas12a has been the subject of extensive investigation both in vitro and in genome editing experiments. However, in vitro studies have often been performed at high magnesium ion concentrations that are inconsistent with the free Mg^2+^ concentrations that would be present in cells. By profiling the specificity of Cas12a orthologs at a range of Mg^2+^ concentrations, we find that Cas12a switches its specificity depending on metal ion concentration. Lowering Mg^2+^ concentration decreases cleavage defects caused by seed mismatches, while increasing the defects caused by PAM-distal mismatches. We show that Cas12a can bind seed mutant targets more rapidly at low Mg^2+^ concentrations, resulting in faster cleavage. In contrast, PAM-distal mismatches cause substantial defects in cleavage following formation of the Cas12a-target complex at low Mg^2+^ concentrations. We observe differences in Cas12a specificity switching between three orthologs that results in variations in the routes of phage escape from Cas12a-mediated immunity. Overall, our results reveal the importance of physiological metal ion conditions on the specificity of Cas effectors that are used in different cellular environments.

## Introduction

Cas12a (formerly Cpf1) is the effector protein of type V-A CRISPR-Cas (clustered regularly interspaced short palindromic repeats-CRISPR associated) immune systems and has been adopted for biotechnology (Murugan et al., 2017; Paul and Montoya, 2020; Zetsche et al., 2015). Cas12a is programmed by a guiding CRISPR RNA (crRNA) to bind and cleave a complementary DNA sequence. In CRISPR-Cas immunity, host cells adapt to infection events by acquiring short segments of the foreign genome within a CRISPR array, which subsequently serves as the template for crRNA production (Lee and Sashital, 2022). In biotechnological experiments, Cas12a can be programmed with a guide RNA of interest designed by the researcher (Paul and Montoya, 2020). In either case, Cas12a binds target DNA based on complementarity between the crRNA and the target strand of the DNA (Zetsche et al., 2015). Following DNA binding, Cas12a uses a metal-ion-dependent RuvC nuclease domain to cleave the non-target strand, followed by a conformational change that exposes the target strand to RuvC cleavage (Cofsky et al., 2020; Son et al., 2021; Stella et al., 2018; Swarts and Jinek, 2019; Swarts et al., 2017). These two successive cleavage events result in double-strand break formation in the target DNA.

The specificity of Cas12a and other Cas effectors used for biotechnology has been extensively studied due to their potential for off-target effects (Fu et al., 2019; Jones et al., 2021; Kim et al., 2016; Kleinstiver et al., 2016; Murugan et al., 2020; Strohkendl et al., 2018). Cas12a specificity is dictated by two main factors: the complementarity between the crRNA guide sequence and the DNA target, and the presence of a PAM (protospacer adjacent motif) next to that DNA target (Zetsche et al., 2015). DNA-targeting Cas effectors like Cas12a require PAM sequences to facilitate target searching and dsDNA destabilization en route to target binding via crRNA-DNA hybridization (Sashital et al., 2012; Semenova et al., 2011; Singh et al., 2018; Sternberg et al., 2014). Following PAM recognition, the crRNA base pairs with the target strand of the DNA, with base pairs in the PAM-proximal “seed” region forming first and base pairs in the PAM-distal region forming last. Mutations in the PAM and seed cause an outsized effect on the ability of Cas12a to bind to a DNA target, resulting in substantial cleavage defects (Kim et al., 2016; Kleinstiver et al., 2016; Zetsche et al., 2015). However, Cas12a cleavage can tolerate mismatches between the crRNA and target DNA, especially when these mismatches occur outside of the seed (Fu et al., 2019; Jones et al., 2021; Murugan et al., 2020; Strohkendl et al., 2018). Mismatches in the PAM-distal region often result in incomplete cleavage of the DNA, where only one strand is cleaved, generating a nicked product (Fu et al., 2019; Murugan et al., 2020).

The differential effects of mismatch location have consequences on the evolution of bacteriophages that are subject to CRISPR-Cas immunity. When targeted in essential genomic regions, phages primarily evade Cas12a and other DNA-targeting Cas effectors by developing mutations in the PAM or seed (Deveau et al., 2008; Schelling et al., 2023; Semenova et al., 2011). However, we recently observed that PAM-distal mutants preferentially emerge when a pre-existing mismatch between the crRNA and target is present in the PAM-distal region (Schelling et al., 2023). This result was surprising, given that previous *in vitro* studies have shown that Cas12a is tolerant of multiple PAM-distal mutations (Fu et al., 2019; Jones et al., 2021; Murugan et al., 2020). The discrepancy between these previous *in vitro* studies and our phage escape data suggested that Cas12a specificity may be altered under physiological conditions. In particular, the concentration of free magnesium ions is expected to be lower in bacterial cells (≤1 mM) than is typically used in *in vitro* studies (5-10 mM) (Froschauer et al., 2004; Tyrrell et al., 2013).

To address this discrepancy, we profiled the specificity of three well-characterized Cas12a orthologs from *Francisella novicida* (FnCas12a), *Acidaminococcus sp.* (AsCas12a), and *Lachnospiraceae bacterium* (LbCas12a) at a range of Mg^2+^ concentrations. Our results reveal a striking specificity switch at low, physiologically-relevant Mg^2+^ concentration. PAM and seed mutants become more tolerated at lower Mg^2+^ concentration, while PAM-distal mutants become less tolerated. Improved cleavage of seed mutant targets at low metal ion concentrations is due to increased rates of DNA binding by Cas12a. In contrast, low metal ion concentration impairs the ability of Cas12a to cleave DNA containing PAM-distal mismatches following target binding. Although we observe the specificity switch for all three orthologs, our results reveal substantial differences in the effects of seed and PAM-distal mismatches on the cleavage activity of AsCas12a in comparison to Lb and FnCas12a. These differences result in marked variability of escaped phage populations that emerge when challenged by each of the three Cas12a orthologs, including the emergence of single PAM-distal escape mutants. Together, our data reveal how alterations to Cas12a specificity under physiological metal ion conditions can determine phage escape outcomes.

## Results

### Mg^2+^ concentration affects off-target and collateral Cas12a cleavage

Although phages are mainly thought to evade CRISPR-mediated immunity by developing PAM and seed mutations (Deveau et al., 2008; Semenova et al., 2011), we previously observed that multiple mismatches in the PAM-distal region can allow for preferential phage escape from Cas12a over seed mutants (Schelling et al., 2023) (Fig. 1A). In head-to-head competition assays, phage strains containing seed or PAM-distal mutations in the Cas12a target region emerge in different distributions depending on the crRNA (Fig. 1A-B). While the seed mutant fully dominates the population when challenged with a perfectly matching crRNA, the distal mutant begins to dominate the population when challenged with a distally mismatched crRNA (Fig. 1B). These results suggest that two PAM-distal mismatches are more deleterious to Cas12a cleavage than the combination of a seed and PAM-distal mismatch, contrary to prior specificity studies of Cas12a (Jones et al., 2021; Kim et al., 2016; Kleinstiver et al., 2016; Murugan et al., 2020). Cleavage analysis of each mutant target using the distally mismatched crRNA revealed the potential source of this discrepancy. While the seed mutant was cleaved much more slowly than the distal mutant at MgCl_2_ concentrations that are typically used for in vitro Cas effector cleavage (10 mM), the distal mutant was cleaved much more slowly than the seed mutant at lower MgCl_2_ concentrations that are more consistent with the concentration present in bacterial cells (1 mM) (Fig. 1C, S1A, B). Surprisingly, the seed mutant target was cleaved ∼5 times faster at low MgCl_2_, while the distal mutant target was cleaved ∼38 times slower.

**Figure 1:**
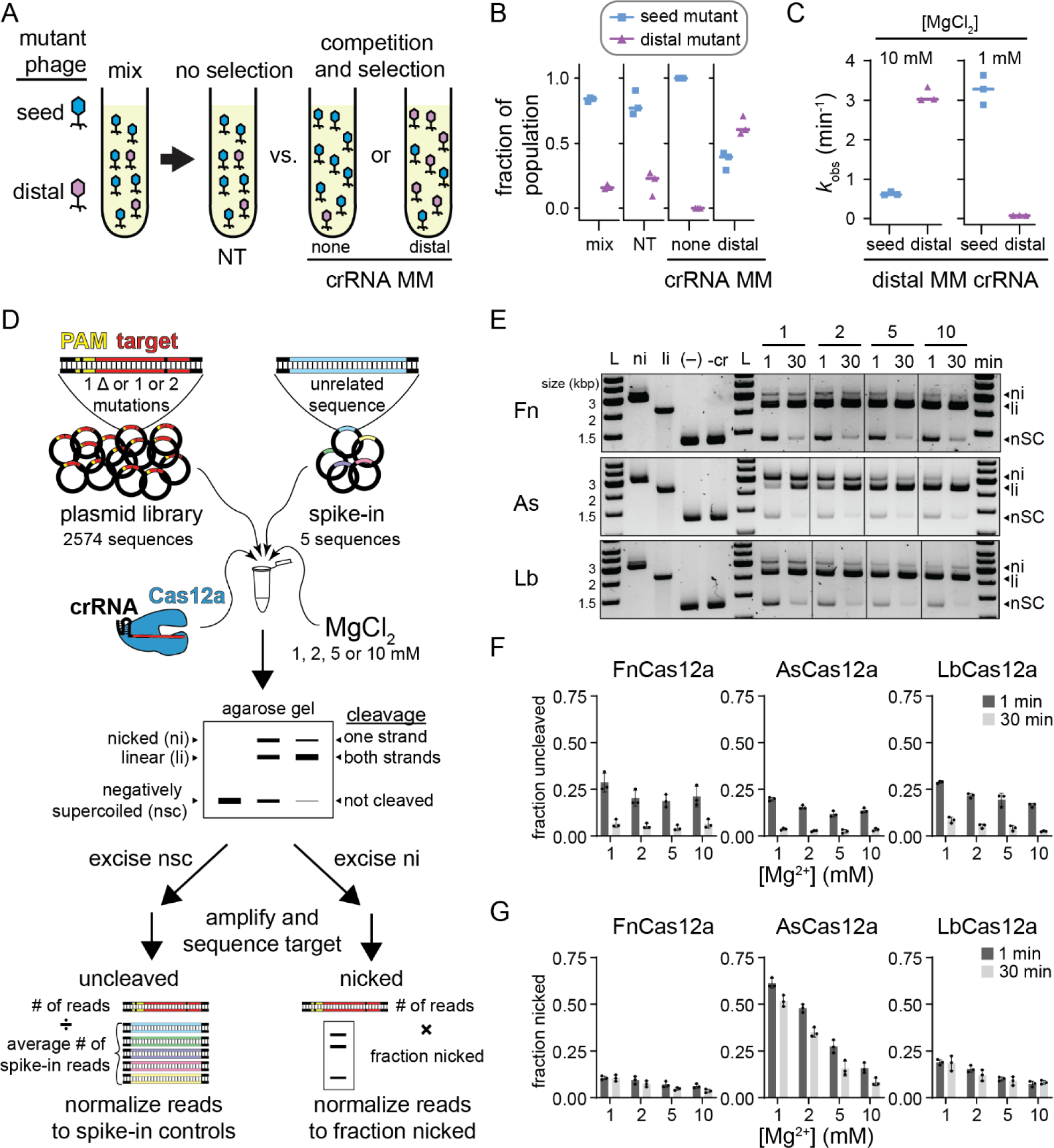
Mg^2+^-dependent Cas12a specificity profiling using plasmid library cleavage (Related to Figure S1 and S2) A) Schematic of phage challenge assay. An initial mix of 80% seed mutant and 20% distal mutant phage were challenged with Cas12a bearing a non-targeting crRNA (no selection), a crRNA matching the original target sequence, or a crRNA containing a mutation at position 15. The two targeting crRNAs result in selection of phage mutants that are more deleterious to Cas12a cleavage. B) Fraction of A2T seed mutant (blue squares) or G17T PAM-distal mutant (purple triangles) in the initial mix, upon challenge by Cas12a bearing a non-targeting crRNA (NT), a crRNA containing no mutations (none), or a crRNA containing mismatch at position 15 (distal). Individual data points from three replicates are shown, and lines represent median values. C) Observed rate constants for cleavage of the A2T or G17T mutant targets with FnCas12a bearing a crRNA containing a mismatch at position 15 in the presence of 10 mM or 1 mM MgCl_2_. Individual data points from three replicates are shown, and lines represent median values. D) Schematic of the plasmid library cleavage assay. Libraries contained all target sequences with single nucleotide deletions or one or two mutations across the PAM and target sequence. Five unrelated sequences were spiked into the sample for normalization. This library was subject to cleavage by Cas12a at four Mg^2+^ concentrations. The cleavage reactions were separated by agarose gel electrophoresis. Negatively supercoiled and nicked bands were excised from the gel, PCR amplified, sequenced, and analyzed as described in Methods. E) Gene L plasmid library cleavage at four Mg^2+^ concentrations for FnCas12a (Fn), AsCas12a (As), and LbCas12a (Lb). ni = nicked, li = linear, nSC = negatively supercoiled. (–) lane contains no protein, -cr contains protein but no crRNA. Both controls contained 10 mM MgCl_2_ and were incubated at 37 °C for 30 min. Gel is representative of three replicates. F-G) Quantification of fraction uncleaved (F) or nicked (G) for the gene L library. The average of three replicates is plotted, with individual data points shown as dots and error bars representing standard deviation.

These results prompted us to re-examine the specificity of Cas12a at a range of Mg^2+^ concentrations. To do this, we used our previously developed *in vitro* plasmid library cleavage assay (Fig. 1D) (Murugan et al., 2020, 2021). In this assay, a Cas12a-crRNA RNA-protein (RNP) complex is used to cleave a negatively supercoiled plasmid library containing target sequences with mutations that introduce mismatches with the crRNA or suboptimal PAM sequences. Following quenching of cleavage, the plasmid library is analyzed by agarose gel electrophoresis allowing separation of uncleaved (negatively supercoiled), nicked (relaxed circular), and fully cleaved (linear) sequences. The uncleaved and nicked fractions of the library are extracted from the gel and subjected to PCR to amplify the target region. These amplicons are then sequenced using MiSeq Illumina sequencing to determine which sequences remained uncleaved and which were cleaved only on one strand.

We selected two targets derived from gene L and gene W of *E. coli* λ phage. We did not observe any major differences in cleavage at 1 and 10 mM MgCl_2_ by three Cas12a orthologs for these targets (Fig. S1C, D). For each target, we created a plasmid library containing all sequences that introduce single-nucleotide deletions or one or two mutations in the PAM or target region and five “spike-in” sequences with unrelated targets as normalization controls for the uncleaved fraction (Fig. 1D, see Methods). We subjected these plasmid libraries to cleavage by Fn, As and LbCas12a at four Mg^2+^ concentrations (1, 2, 5 and 10 mM), collecting two time points at 1 and 30 min (Fig. 1E-G, S1E-G). We observed that ∼15-30% of the library remained uncleaved at 1 min and that most of the library was cleaved by 30 min (Fig. 1F, S1F). Variations in the fraction uncleaved at different Mg^2+^ concentrations were relatively minor. In contrast, we observed a substantial decrease in the amount of nicked DNA as Mg^2+^ concentration increased, especially for AsCas12a (Fig. 1G, S1G). Interestingly, the amount of nicked DNA remained similar between the 1 min and 30 min time points for FnCas12a and LbCas12a, suggesting that some sequences are rapidly nicked by these orthologs, but are never fully cleaved. Nicking of the plasmid libraries suggests that AsCas12a has a metal-ion-dependent second-strand cleavage defect that is more pronounced than for Fn and LbCas12a.

Notably, Cas12a also displays “collateral” non-specific DNA degradation activity upon activation by a target sequence (Chen et al., 2018; Li et al., 2018). We previously observed non-specific nicking of plasmids by both Fn and LbCas12a when activated with a perfectly matching or mismatched target (Murugan et al., 2020). However, it is unclear whether non-specific cleavage occurs at lower Mg^2+^ concentration. We tested whether plasmids containing the perfect gene L and W target could activate LbCas12a for cleavage of an empty plasmid at the four metal ion concentrations used for plasmid library cleavage (Fig. S2A). We observed substantially more nicked DNA at higher Mg^2+^ concentrations, but very little at 1 mM Mg^2+^, suggesting that activated nicking is metal ion dependent. Similarly, both L and W target plasmid libraries could activate LbCas12a, and to a lesser degree FnCas12a, for cleavage of an empty plasmid, with the most nicking observed at the highest Mg^2+^ concentration (Fig. S2B). Spike-in plasmids in the plasmid library allowed us to account for non-specific nicking of sequences that cannot otherwise be cleaved by Cas12a (Fig. 1D, see Methods).

We also assessed the ability of all three orthologs to degrade single-stranded DNA upon activation (Fig. S2C). Interestingly, Fn and AsCas12a did not degrade ssDNA at 1 and 2 mM Mg^2+^, while LbCas12a degraded ssDNA more slowly at lower Mg^2+^ concentrations. These results may explain why Cas12a non-specific ssDNA degradation activity was recently shown to have no effect on phage defense (Marino et al., 2022), given that this activity is decreased at physiological metal ion concentrations.

### PAM-proximal and distal mutants are differentially affected by Mg^2+^ concentration

Following cleavage of the plasmid library, we next isolated and PCR amplified the uncleaved and nicked fractions and subjected the amplicons to high-throughput sequencing (Fig. 1D). We determined fraction uncleaved and nicked values for sequences present in each sample (see Methods). To rapidly visualize the effects of MgCl_2_ concentration on mutant sequence cleavage, we created volcano plots comparing the fraction uncleaved or nicked at 1 and 10 mM MgCl_2_ (Fig. 2A-B, S3). Sequences with large and statistically significant changes were colored based on the location of the first mutation in the sequence for the uncleaved fraction (Fig. 2A, S3A) or the location of the second mutation for the nicked fraction (Fig. 2B, S3B). These plots reveal that sequences with the first mutation in the PAM or seed are more likely to be uncleaved at higher MgCl_2_ concentration, especially for Fn and LbCas12a following 30 min of cleavage (Fig. 2A, S3A). In contrast, sequences with the first or second mutation in the mid or PAM-distal region are more likely to be uncleaved or nicked at lower MgCl_2_ concentration (Fig. 2A-B, S3A-B). Notably, although we observed many sequences where the first mutation was in the PAM or seed region that were relatively enriched in the uncleaved fraction at lower MgCl_2_ concentration, most of these sequences had a second mutation in the mid or PAM-distal region (see below).

**Figure 2:**
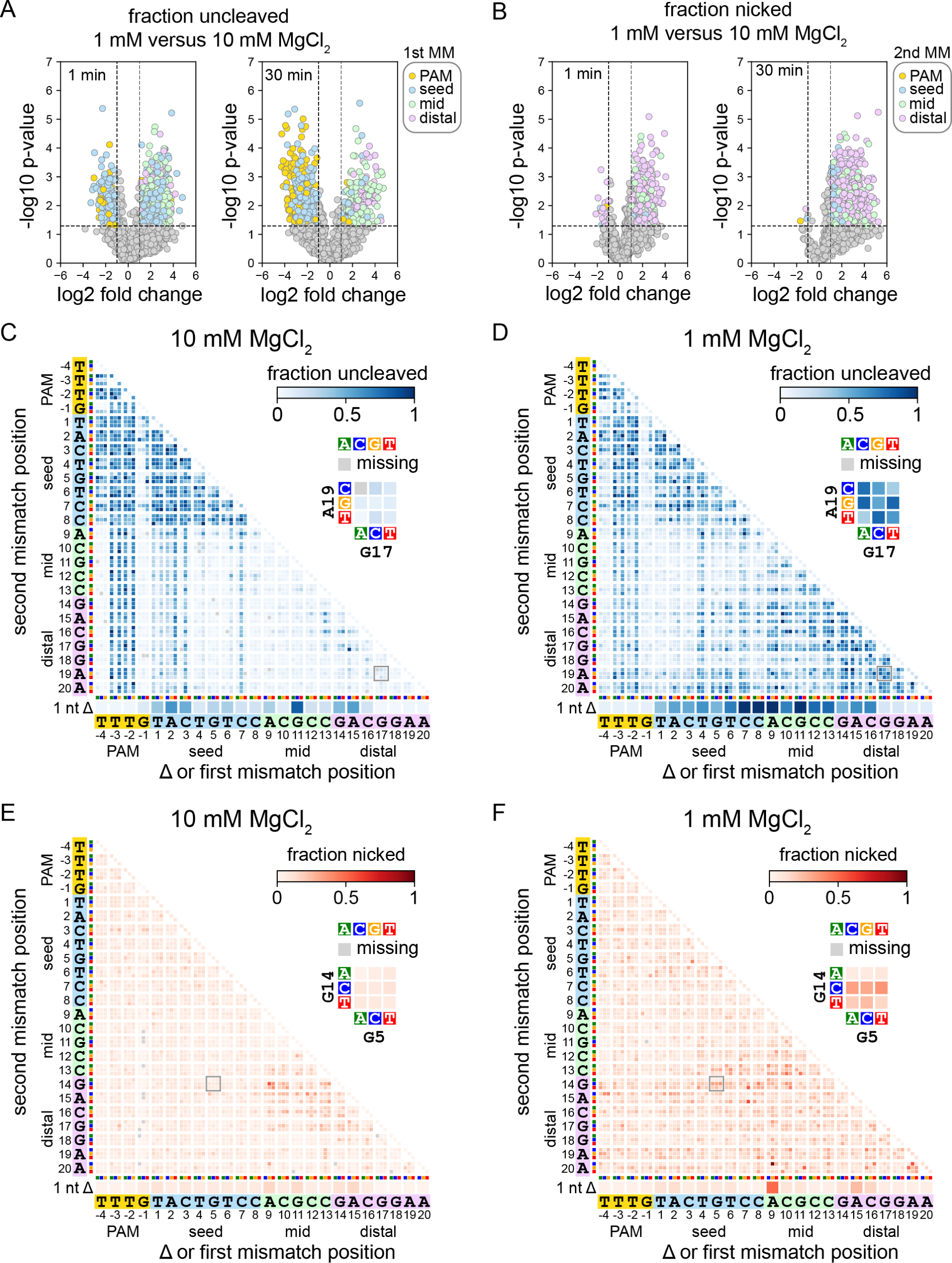
Specificity is altered in both the seed and the PAM-distal region at lower Mg^2+^ concentration (related to Figure S3 and Supplementary Data 1) A-B) Volcano plots comparing sequences present in uncleaved (A) or nicked (B) fractions following cleavage of the gene L library at 1 or 10 mM MgCl_2_ by FnCas12a for 1 or 30 min. Data points in (A) are colored by the location of the first mutation in the sequence. Data points in (B) are colored by the location of the second mutation in the sequence. *P* values compare up to three replicates using an unpaired two-tailed *t* test C-D) Heatmaps showing the abundance of mutant sequences from the gene L plasmid library in the uncleaved (negatively supercoiled) fraction following 1 min cleavage by FnCas12a in the presence of 10 mM (C) or 1 mM (D) MgCl_2_. E-F) Heatmaps showing the abundance of mutant sequences from the gene L plasmid library in the nicked fraction following 1 min cleavage by FnCas12a in the presence of 10 mM (E) or 1 mM (F) MgCl_2_. Sequences with a single nucleotide deletion are shown at the bottom of each heatmap. Sequences with one mutation are shown along the diagonal of the 2D heatmaps. Sequences with mutations at two positions are depicted as 3×3 arrays where each box represents one combination of mutations at the two positions. For each heatmap, one 3×3 array is shown in close-up. Missing sequences are depicted in the heatmap with a gray box.

To more thoroughly investigate which sequences were uncleaved or nicked at each MgCl_2_ concentration, we plotted heatmaps that enable visualization of the abundance of each mutant sequence in each fraction (Jones et al., 2021; Jung et al., 2017) (Fig. 2C-F, Supplementary Data 1). These heatmaps reveal the exact types of mutations that cause cleavage defects at each Mg^2+^ concentration. For example, for FnCas12a cleaving the gene L library, we observed that single transversion mutations in the first three positions of the seed caused an enrichment of these sequences at 10 mM MgCl_2_ (Fig 2C, diagonal of the heatmap). These seed mutations in combination with mutations in the mid or PAM-distal region of the target were also enriched at high Mg^2+^. In contrast, sequences with individual point mutations at the first three positions of the seed, or combinations of these seed mutations with mid or PAM-distal mutations were depleted from the uncleaved fraction at 1 mM MgCl_2_ (Fig. 2D). Instead, we observed a substantial increase in uncleaved sequences containing two mismatches in the mid and/or PAM-distal region at low Mg^2+^.

The nicked fraction likely includes mutated sequences that can undergo cleavage on the non-target strand, but do not undergo the second cleavage step on the target strand (Fu et al., 2019; Murugan et al., 2020; Swarts and Jinek, 2019). Plotting the fraction nicked on heatmaps revealed that nicking of the gene L library by FnCas12a is largely caused by combinations of mutations at positions 9-17 of the target at 10 mM MgCl_2_ (Fig. 2E), similar to previous reports that PAM-distal mutations cause incomplete cleavage by Cas12a (Fu et al., 2019; Murugan et al., 2020). At 1 mM MgCl_2_, double mutations in which one mutation was in the mid or distal region were relatively enriched in the nicked fraction (Fig. 2F), although sequences that were highly enriched in the uncleaved fraction were mostly depleted from the nicked fraction (Fig. 2D, F). These results suggest that mutations that cause incomplete cleavage at higher divalent metal ion concentrations may cause a complete loss of cleavage at lower ionic conditions.

Overall, our data yielded 16 heatmaps each for the uncleaved and nicked fractions per ortholog (4 Mg^2+^ concentrations for two timepoints and two libraries). To simplify viewing of these heatmaps, we created animations for each timepoint and library, in which each frame shows the uncleaved or nicked fraction heatmap for a given Cas12a ortholog at each Mg^2+^ concentration (Supplementary Data 1). The animations for the uncleaved fraction reveal similar trends in the PAM-distal region for all orthologs and both libraries, where several sequences with deletions or mutations in the PAM-distal region are highly abundant in the uncleaved fraction at lower Mg^2+^ concentrations, even after 30 min of cleavage. The improved tolerance of seed mutations at lower Mg^2+^ was also most pronounced FnCas12a after 30 min cleavage. Double seed mutants or sequences containing combinations of PAM and mid/distal mutations were depleted from the uncleaved fraction at lower Mg^2+^ concentrations for both libraries, suggesting a higher tolerance for mutations in both the PAM and seed by FnCas12a at low Mg^2+^. Similar tolerance of double mutants containing seed and PAM mutants was less pronounced at lower Mg^2+^ for As and LbCas12a following 30 min of cleavage, suggesting that Mg^2+^-dependent PAM/seed mutation tolerance varies between different Cas12a orthologs.

For the nicked fraction, we observed similar accumulation of nicked sequences with at least one mismatch in the mid or distal region at low Mg^2+^, with AsCas12a showing the strongest second strand cleavage defect. At high Mg^2+^ concentration, nicked sequences mostly localize to a region of the heatmap representing dual mutations in the mid and PAM-distal region, while at low Mg2+, combinations of mutations in the PAM/seed and mid/distal regions result in substantial nicking. Notably, several of the nicked heatmaps are similar between the two time points, suggesting that sequences that were nicked at the early timepoint remained nicked throughout the course of the experiment.

### Visualizing the metal-dependent specificity switch

In our plasmid library cleavage assay, the linear DNA fraction that has been fully cleaved by Cas12a cannot be PCR amplified due to cleavage of both strands of the potential PCR template. However, our normalization strategy to determine what fraction of individual sequences remained uncleaved or were nicked allowed us to estimate what fraction of the DNA was fully cleaved (see Methods). Sequences that were highly abundant in either the uncleaved fraction, the nicked fraction, or both, were not fully cleaved, while sequences that were depleted from both the uncleaved and nicked fraction were fully cleaved (Fig. 3A, B). Determining the fraction of DNA that was fully cleaved allowed us to calculate slope values that report the change in cleavage of individual sequences in the plasmid library across the four Mg^2+^ concentrations. Negative slope values indicate sequences that decreased in cleavage as Mg^2+^ increased (Fig. 3A), while positive slope values indicate sequences that increased in cleavage as Mg^2+^ increased (Fig. 3B).

**Figure 3:**
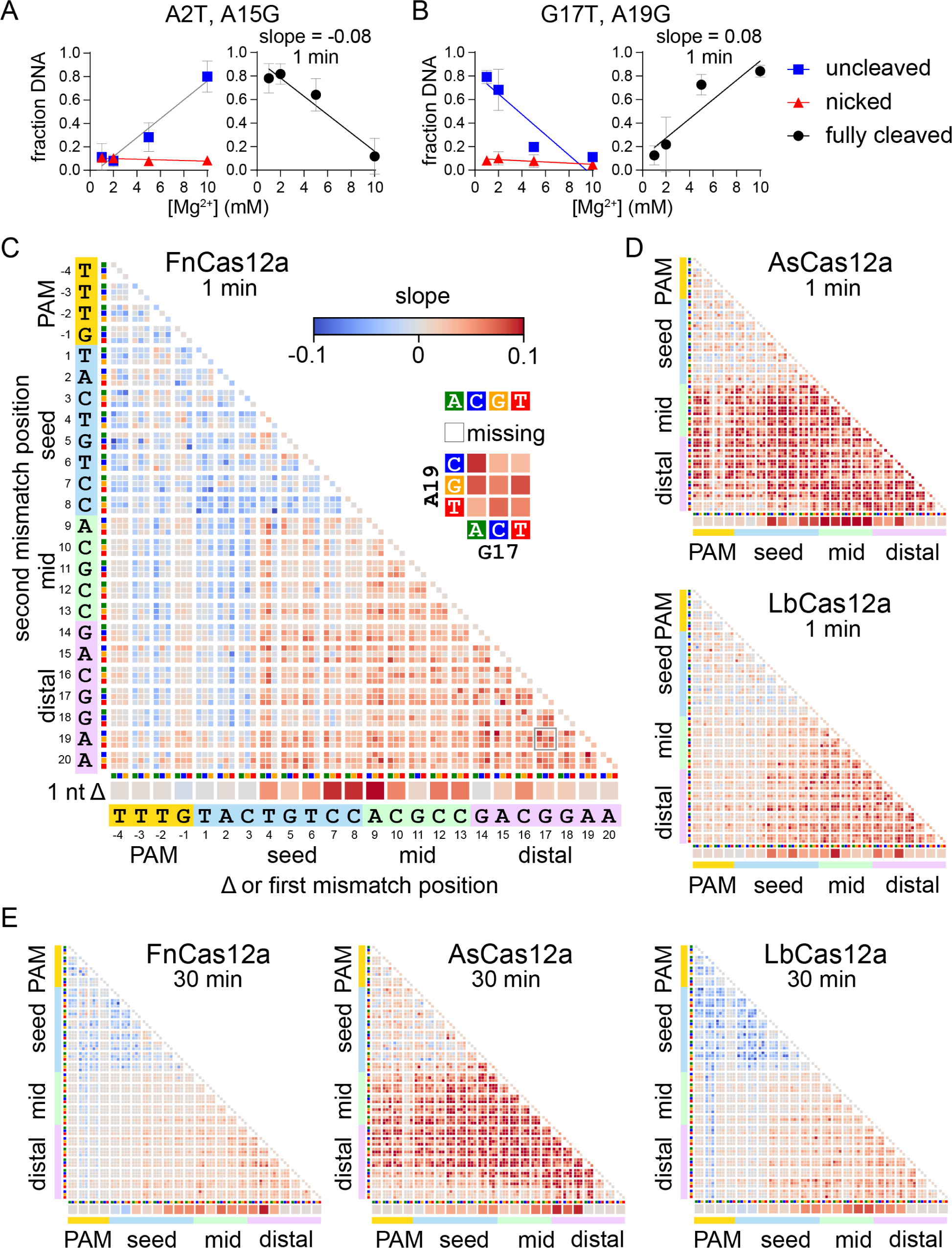
Metal-dependent specificity switching for three Cas12a orthologs (Related to Figure S4) A-B) Cleavage is linearly related to Mg^2+^ concentration. The abundance of gene L A2T A15G (A) or G17T A19G (B) in the uncleaved (negatively supercoiled, blue boxes) or nicked (red triangles) fraction is plotted versus Mg^2+^ concentration and each curve is fit to a linear regression. These values were used to determine the fraction of DNA that was cleaved on both strands to produce linear, fully cleaved DNA (see Methods), which is plotted on the right and fit to a linear regression. The average of three replicates is plotted, with error bars representing standard deviation. C) Heatmap as in Figure 2 plotting slopes for fully cleaved DNA versus Mg^2+^ as determined in panels (A-B). The heatmap is for the gene L target plasmid library following 1 min cleavage by FnCas12a. Missing sequences are represented by white boxes. D) Slope heatmaps of the gene L target plasmid library following 1 min cleavage by AsCas12a or LbCas12a. E) Slope heatmaps of the gene L target plasmid library following 30 min cleavage by all three orthologs.

We plotted slope values for the change in fully cleaved DNA across Mg^2+^ concentrations in heatmaps similar to those shown in Fig. 2 (Fig. 3C-E, S4). These heatmaps allow for rapid visualization of the different effects that changes in Mg^2+^ concentrations have across the target sequence. For the gene L target, after 1 min of cleavage by FnCas12a, we observed negative slopes for individual transversion mutations in the PAM (positions -3 and -2) and the seed (positions 1-3) as well as for double mutants containing one of these initial mutations (Fig. 3C). In addition, most double mutations in the PAM and seed had negative slopes. Conversely, sequences with deletions or double mutations at positions 4-20 mostly have positive slopes (Fig. 3C). While we observed similar trends in the PAM-distal region for LbCas12a, the heatmap for AsCas12a is strikingly different, with almost all sequences containing at least one mutation in the mid or PAM-distal region having large positive slopes (Fig. 3D). This difference reflects the significant defect in second strand cleavage observed at low Mg^2+^ concentrations for As, but not for Fn or LbCas12a (Fig. 1G). Positive slopes were less pronounced at 1 min for As and LbCas12a than for FnCas12a, and limited to a few double mutants in the PAM and seed (Fig. 3D). However, at 30 min, we observed substantially more negative slopes for mutants in the PAM and seed when cleaved by LbCas12a (Fig. 3E).

For the gene W target, we observed the most pronounced negative slopes for sequences containing two mutations in the PAM and seed for both FnCas12a at both time points, and to a lesser degree for As and LbCas12a (Fig. S4). For FnCas12a, we also observed slightly negative slopes for some sequences in which single nucleotides in the first few positions of the seed were deleted. We again observed positive slopes for sequences containing at least one mutation in the mid or PAM-distal region for all three orthologs. Overall, these data suggest that Mg^2+^ concentration has consistent effects on Cas12a ortholog specificity for different targets, with some variations depending on target sequence.

### Varied mechanisms of Mg^2+^-dependent specificity switching

Our plasmid library cleavage results reveal that variations in Mg^2+^ concentration do not uniformly affect mismatch tolerance across the target region. PAM mutations and seed mismatches are more tolerated at low Mg^2+^ concentration, especially for FnCas12a. In contrast, mismatches in the mid or distal region are less tolerated at low Mg^2+^ concentration. We next sought to determine what step of the Cas12a target binding and cleavage cycle is most affected by lower Mg^2+^ concentrations for sequences containing seed or PAM-distal mismatches. We chose to further study FnCas12a, which had the most pronounced specificity switch for both seed and PAM-distal mismatched sequences. We measured the rate of cleavage for individual target sequences by initiating the cleavage reaction in two different ways (Fig. 4A). Following Cas12a RNP formation, the RNP was either mixed with DNA in the presence of Mg^2+^ (initiation with DNA) or the RNP and DNA were mixed and incubated for 30 min prior to the addition of Mg^2+^ (initiation with Mg^2+^). When the reaction is initiated by addition of DNA, the rate of nicked or linear product formation is dependent on the rate of both binding and the first or second cleavage event, respectively. In contrast, initiation with Mg^2+^ enables measurement of the rate of the two cleavage steps following RNP-DNA binding.

**Figure 4:**
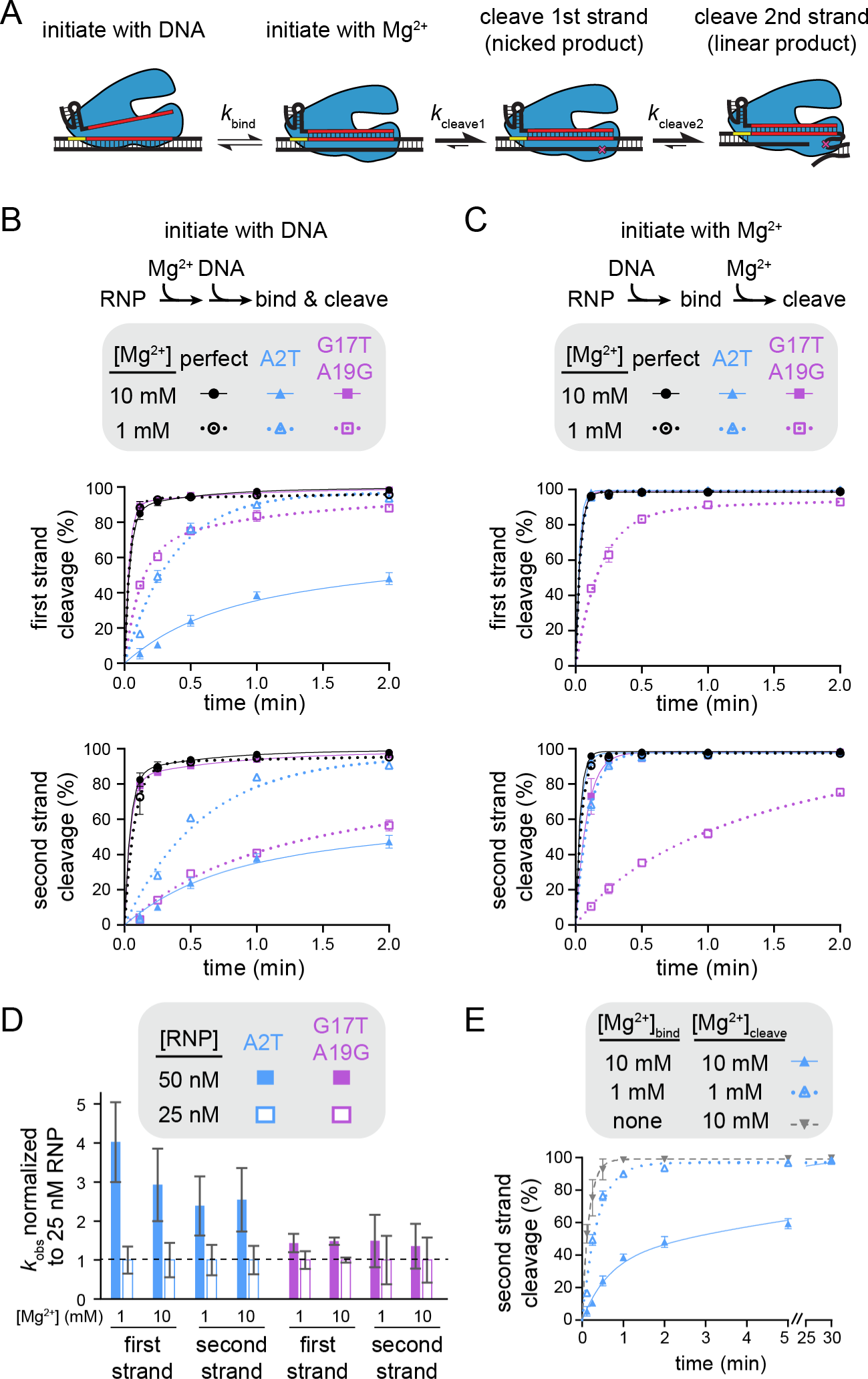
Mechanism of specificity switching varies for seed and PAM-distal mutants (Related to Figure S5) A) Schematic representing different potential rate-limiting steps during Cas12a catalysis. The rates of binding or each cleavage step is represented with a different forward rate constant. B) Cleavage of gene L targets by FnCas12a in which cleavage was initiated by mixing Cas12a RNP with DNA. The gene L target containing a perfect match (black), an A2T seed mutation (blue), or a G17T A19G PAM-distal double mutant (purple) were cleaved by FnCas12a in the presence of 10 mM Mg^2+^ (solid shapes and lines) or 1 mM Mg^2+^ (open shapes and dotted lines). The rate of cleavage of the first strand and the second strand are plotted separately (see Methods). The average of three replicates is shown and error bars represent standard deviation. C) Cleavage of gene L targets by FnCas12a in which RNP and DNA were incubated in the absence of Mg^2+^ prior to initiation of cleavage through the addition of Mg^2+^, as in (B). D) Normalized rate constants derived from cleavage of gene L A2T mutant (blue) or G17T A19G mutant (purple) at 50 nM (solid bars) or 25 nM (white outlined bars) FnCas12a RNP concentrations. For each condition, the average rate constant values (n = 3 or 4) at each concentration were normalized to the 25 nM rate constant value and standard deviation was propagated for division. E) Cleavage of the A2T gene L mutant target by FnCas12a in the presence of 10 mM Mg^2+^ (solid triangle and lines, blue), 1 mM Mg^2+^ (open triangle and dotted lines, blue), or no Mg^2+^ (upside down triangle and dashed line, gray) during RNP-DNA binding. The average of three replicates is shown and error bars represent standard deviation.

We selected a seed mutant (A2T) and a dual PAM-distal mutant (G17T A19G) for further cleavage analysis. As expected, we observed faster cleavage of the seed mutant and slower cleavage of the PAM-distal mutant at 1 mM Mg^2+^ in comparison to 10 mM Mg^2+^ when cleavage was initiated with DNA (Fig. 4B, S5A). While the first cleavage event occurred at a similar rate for the two mutants at 1 mM Mg^2+^, the PAM-distal mutant target was linearized slowly after initial nicking (Fig. S5A), resulting in a much slower second strand cleavage rate for the distal mutant in comparison to the seed mutant at the lower Mg^2+^ concentration (Fig. 4B, bottom).

When the DNA targets were pre-bound to Cas12a and cleavage was initiated by adding Mg^2+^, first-strand cleavage of the perfectly matching and the seed mutant targets was complete by the first time point (7 s) at both Mg^2+^ concentrations, as well as for the PAM-distal mutant at 10 mM Mg^2+^ (Fig. 4C, S5B). These results indicate that all three DNA constructs were fully bound to the Cas12a RNP prior to initiation of cleavage through the addition of Mg^2+^, and that any differences in the rate of first-strand cleavage following DNA binding are unmeasurable. Thus, first-strand cleavage defects for the seed mutant target observed upon initiation with DNA can be mainly attributed to DNA binding defects, which are exacerbated at higher Mg^2+^ concentration.

In contrast, we observed marked decreases in the rates of both first- and second-strand cleavage of the G17T A19G mutant when cleavage was initiated with 1 mM Mg^2+^ (Fig. 4C, S5B). Indeed, the rate of cleavage of this target was similar regardless of the method of cleavage initiation (Fig. S4A-B, bottom right gel of each panel). The similar cleavage defect for the pre-bound Cas12a-target complex suggests that DNA binding is not rate determining for Cas12a cleavage of the PAM-distal mutant target at low Mg^2+^ concentration. Instead, steps that occur following DNA binding but prior to cleavage of each strand are slowed at low Mg^2+^, most likely conformational changes that are required to position each strand into the RuvC active site (Son et al., 2021). Consistently, while cleavage rate was dependent on Cas12a concentration for the seed mutant target, we did not observe a concentration-dependent change in cleavage rate of the PAM-distal mutant target (Fig. 4D, S5A, S5C).

The observation that a higher Mg^2+^ concentration inhibits Cas12a binding to the seed mutant target is surprising, given that divalent metal ions could electrostatically stabilize protein-nucleic acid interactions. We reasoned that Mg^2+^-dependent stabilization of the DNA duplex may inhibit DNA unwinding by Cas12a when a seed mismatch is present (Owczarzy et al., 2008). Indeed, when RNP-DNA binding was initiated in the absence of Mg^2+^ prior to addition of Mg^2+^ at various time points (see Methods), cleavage of the A2T target was even faster than at 1 mM Mg^2+^ (Fig. 4E, S5D). Together, these results suggest that Cas12a seed-mutant target binding is inhibited at higher Mg^2+^ concentrations due to stronger DNA unwinding defects.

### Phage escape outcomes reveal consequences of Cas12a ortholog specificities

Our specificity profiling revealed differences in the degree of mismatch tolerance for the three Cas12a orthologs at low Mg^2+^ concentration (Fig. 3C-F, S4). In particular, for AsCas12a, low Mg^2+^ concentration had less of an effect on cleavage of seed mutant sequences, but caused more deleterious second-strand cleavage defects than for Fn and LbCas12a. We next investigated whether these differences in specificity impact what types of mutant phages emerge when challenged by each Cas12a variant. We created a panel of 8 crRNAs targeting different regions of the λ_vir_ genome and co-expressed each combination of Cas12a ortholog and crRNA in *E. coli* K12. Two of these targets were located just upstream or downstream of coding regions (gene J and gene D, respectively), while the remainder of the targets were located within the coding region of essential genes. Cas12a was expressed using a weak promoter (Phan et al., 2019; Schelling et al., 2023) to maximize the likelihood that mutant phages might escape targeting and become abundant in the population. After infecting triplicate cultures with λ_vir_, we sampled the phage population at various time points and PCR amplified the target region of the phage genome. These amplicons were then subjected to high-throughput sequencing to determine whether and which mutants were present in the population following Cas12a-mediated defense.

For most AsCas12a and FnCas12a cultures, we observed a period of growth followed by delayed lysis resulting in a low culture density at 12 hr, which is characteristic of emergence of escape mutants in the phage population (Fig. 5A, S6A) (Schelling et al., 2023). However, we did not observe lysis for at least one replicate of most LbCas12a cultures. These differences may be due to relatively high expression of LbCas12a, as determined by Western blots of cultures expressing the three Cas12a orthologs (Fig. S6B). Lysis was generally accompanied by nearly complete mutagenesis of the phage population (Fig. 5B). All phage lysates from AsCas12a cultures were >98% mutated, as were most of the FnCas12a cultures. For cultures where we did not observe lysis, for example FnCas12a gene C, we generally observed incomplete or no mutagenesis, as was the case for three LbCas12a cultures that did not lyse (genes C, E, and W).

**Figure 5:**
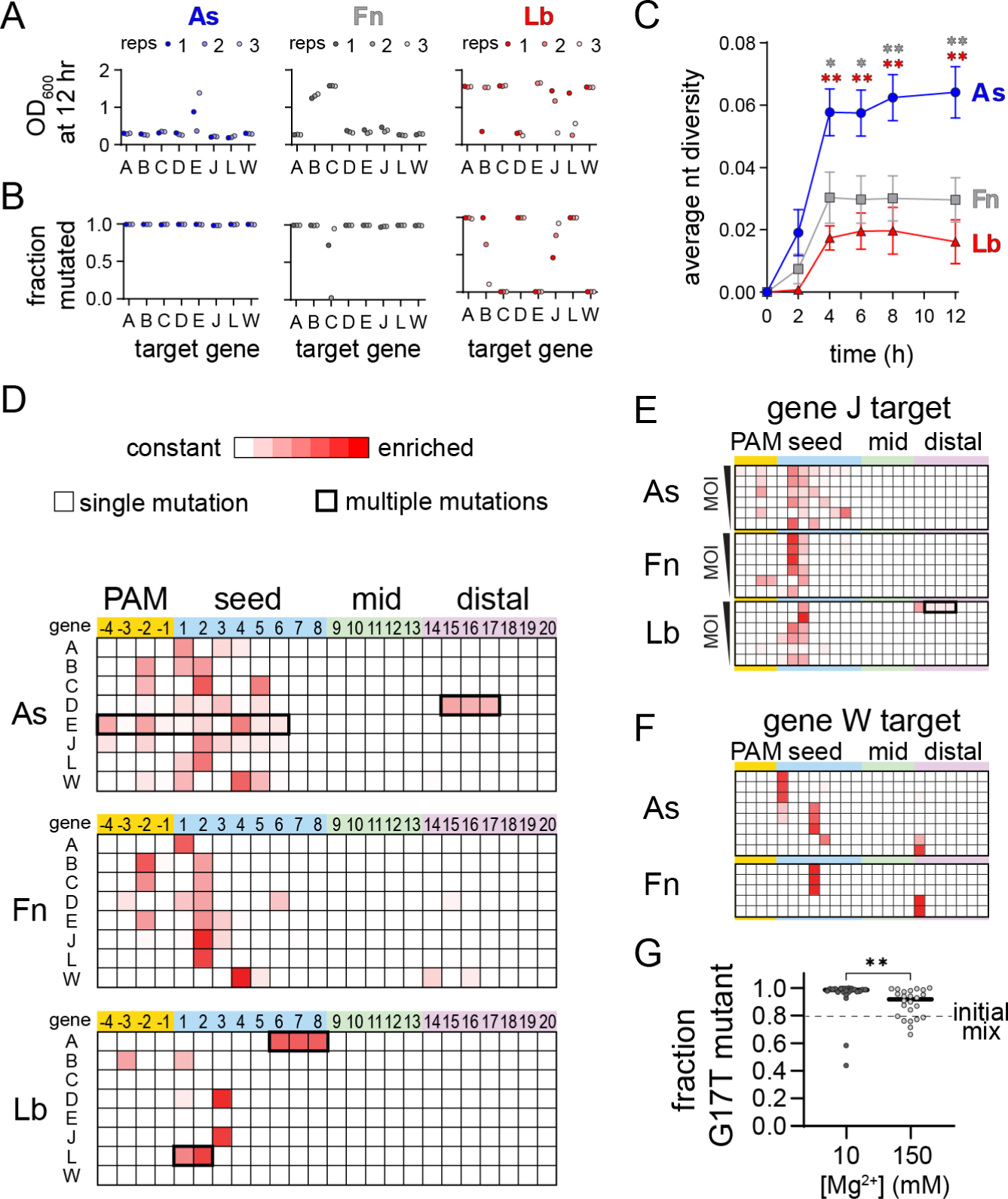
Cas12a orthologs have varied escape outcomes (Related to Figure S6) A) Optical density at 600 nm of phage-infected cultures of *E. coli* expressing the indicated Cas12a ortholog and a crRNA targeting the indicated gene following 12 h. Low values indicate full lysis, while values >1 indicate culture survival. The full growth curves are shown in Fig. S5A. B) Fraction of phage population for each culture in (A) in which at least one mutation was present in the PAM or targeted region of the phage genome. C) Average nucleotide diversity over time of all phage populations from cultures in which mutant phages emerged (see Methods). The error bars represent standard error of the mean of the average nucleotide diversity for between 5 to 8 target sequences. * *P* ≤ 0.05, ** *P* ≤ 0.01 as determined by an unpaired two-tailed *t* test comparing scores for AsCas12a and FnCas12a (gray, upper asterisks) or AsCas12a and LbCas12a (red, lower asterisks). D) Locations of mutations present in phage populations isolated from three infected *E. coli* cultures expressing the indicated Cas12a ortholog and a crRNA targeting the indicated λ_vir_ gene following 12 h of infection at an MOI of 0.8. Heatmaps plot Z-score values (scale of 0 to 7.5) for the presence of mutations in phage populations from infected cultures versus an uninfected control. Shades of red indicate the degree of enrichment of mutations at each position of the target. Most mutations were single nucleotide variants (thinner line around box), although multiple mutations arose in some cultures (thicker line around box). E) Locations of mutations present in phage populations isolated from infected *E. coli* cultures expressing the indicated Cas12a ortholog targeting gene J following 8 h of infection. The cultures were infected with phage at an MOI of 1, 0.5, 0.15, 0.08, 0.04, or 0.02 from top to bottom. Z-score are plotted as in (D). F) Locations of mutations present in phage populations isolated from infected *E. coli* cultures expressing the indicated Cas12a ortholog targeting gene W following 8 h of infection an MOI of 0.08. Only replicates in which the cultures underwent lysis are shown (out of 12 total). Z-score are plotted as in (D). G) Outcomes of competition assays between a seed and PAM-distal mutant phage in cultures grown in media supplemented with 10 mM or 150 mM MgSO_4_. The initial mix contained an 80:20 distribution of the G17T and A2T mutants. The fraction of the phage population containing the single G17T point mutation following lysis of the culture is plotted for 24 replicates. The approximate *P* value is 0.002 based on a nonparametric unpaired Kolmogorov-Smirnov test.

We next examined what types of mutations arose within the targeted regions. An overall analysis of the nucleotide diversity present in each phage lysate revealed that challenge by AsCas12a produced the most diverse populations for cultures in which mutant phages emerged (Fig. 5C). Heatmaps plotting the locations of mutations across the PAM and seed underscore the diversity of the mutant populations that escaped from AsCas12a, in comparison to both Fn and LbCas12a (Fig. 5D). Single point mutations emerged throughout the PAM and seed when phages were challenged with AsCas12a, but were mainly localized to the most deleterious positions (-3 and -2 positions of the PAM or positions 1-3 of the seed) for Fn and LbCas12a. Although most mutants contained a single nucleotide substitution, we observed some sequences with multiple mutations, including deletion of consecutive nucleotides (gene D for AsCas12a and gene A for LbCas12a) or multiple point mutations that developed over time (gene E for AsCas12a and gene L for LbCas12a, Fig. S6C, D).

Our initial analysis suggested that a broader range of seed mutations could allow for escape from AsCas12a in comparison to Fn or LbCas12a. It is possible that these differences are due to expression level differences between the three Cas12a variants (Fig. S6B), as less deleterious mutations may be more likely to emerge when less Cas12a is available. To determine whether mutant diversity decreased as a function of Cas12a:target ratio, we performed phage escape assays at a range of multiplicities of infection (MOIs). We selected the gene J target, for which we observed mutations across the PAM and seed region for AsCas12a, but only at positions 2 or 3 of the seed for Fn and LbCas12a (Fig. 5D). We continued to observe diverse mutations across the PAM and seed for AsCas12a across MOIs, while mutations were mainly limited to the most deleterious positions or to PAM-distal deletions for Fn and LbCas12a at most MOIs (Fig. 5E, S6E). These results, along with differences observed in our specificity profiling data, suggest that AsCas12a differentially tolerates seed mutations under physiological conditions in comparison to Fn and LbCas12a.

Our specificity profiling also suggested that PAM-distal mismatches could be highly deleterious to second-strand cleavage by Cas12a at low metal ion concentrations (Fig. 2B, S3, Supplementary Data 1), suggesting that PAM-distal mutants may emerge in phage populations challenged by Cas12a. Indeed, for the gene W target, we observed PAM-distal point mutations for the FnCas12a culture (Fig. 5D). To further test this, we performed an additional 12 phage infection replicates for As and FnCas12a bearing a crRNA against the gene W target. Out of the 12 cultures, 8 lysed for AsCas12a and 5 lysed for FnCas12a. Interestingly, we observed a single point mutation at position 14 for three of these lysed cultures (Fig. 5F). These results suggest that individual PAM-distal mutations can be sufficiently deleterious to Cas12a cleavage to allow for phage escape.

### Metal-dependent specificity can alter phage-escape outcomes

Finally, we wondered whether fluctuations in Mg^2+^ concentrations in bacterial cells may alter the distribution of mutant phages in populations challenged by Cas12a. We repeated a competition assay similar to the experiment shown in Fig. 1A-B, but now growing the *E. coli* cultures in media supplemented with either 10 or 150 mM Mg^2+^ (Fig. 5G). Starting with an initial mix favoring the PAM-distal mutant over the seed (80:20 distribution), we observed that the PAM-distal mutant rose to >95% of the population in almost all cultures (21/24, median value 98.6%) at 10 mM Mg^2+^ when challenged with Cas12a bearing a distally-mismatched crRNA. In contrast, we observed a significantly different distribution of the PAM-distal mutant in cultures grown at 150 mM Mg^2+^ with the PAM-distal mutant rising to >95% of the population in only 9 out of 24 cultures (median value 91.8%). These results are consistent with the reduced impact of PAM-distal mismatches on Cas12a cleavage at higher Mg^2+^ concentration, and suggest that changes in cellular Mg^2+^ concentrations can affect phage escape outcomes.

## Discussion

Cas effector specificity is tuned to meet the many demands of CRISPR-Cas immunity. Cas effectors must rapidly and selectively locate targets in foreign DNA while also allowing some tolerance of mismatches between the crRNA and target to mitigate immune evasion. Here, we show that an additional factor of divalent metal ion availability also tunes the specificity of Cas12a. We find that Cas12a is more tolerant of seed mutations at lower Mg^2+^ concentration. This increased seed mismatch tolerance is offset by a decreased tolerance for mutations in the PAM-distal region. These results explain our observations that PAM-distal mutant phages preferentially emerge upon challenge by Cas12a bearing a distally-mismatched crRNA (Schelling et al., 2023). Together, these results strongly suggest that the specificity profile we measured at lower Mg^2+^ concentration is physiologically relevant and reflects the degree of Cas12a cleavage that would occur against mismatched targets in cells.

Seed mismatches cause binding defects due to directional target unwinding, in which mutations that are close to the PAM are encountered early in R-loop formation and can more readily lead to R-loop collapse (Rutkauskas et al., 2015; Singh et al., 2018; Szczelkun et al., 2014). Consistently, our analysis of Mg^2+^-dependent Cas12a cleavage revealed that binding is rate limiting for a seed mutant target, and that binding defects are more pronounced at higher Mg^2+^ concentration. In the absence of Mg^2+^, Cas12a bound a seed mutant target relatively rapidly. Notably, in our cleavage assays, we used a negatively supercoiled DNA target plasmid. Negative supercoiling has been shown to improve the ability of DNA-binding Cas effectors to their targets, as well as to off-target sites (Newton et al., 2023; Westra et al., 2012). Our results now reveal the additional role of divalent metal ions, which may decrease the ability of Cas effectors to unwind DNA by increasing the stability of the DNA double helix (Owczarzy et al., 2008). Future studies will be required to determine whether metal ion concentration has similar effects on the ability of diverse Cas effectors to unwind both mutated and perfectly matched target sequences.

Our results reveal that at low Mg^2+^ concentrations, cleavage of targets containing multiple PAM-distal mismatches is strongly rate-limited by steps that occur after target binding, especially for the second, target-strand cleavage event. Cas12a requires multiple conformational changes for non-target and target strand cleavage (Naqvi et al., 2022; Saha et al., 2020; Son et al., 2021; Stella et al., 2018), which may be inhibited by PAM-distal mismatches. In particular, the Mg^2+^-dependent conformational change required for target strand cleavage may be substantially inhibited in the presence of PAM-distal mismatches (Son et al., 2021). Alternatively, these defects may be due to R-loop instability in the PAM-distal region. Divalent metal ions may help to stabilize the full R-loop when mismatches are present in the PAM-distal region, preventing unwinding of PAM-distal base pairs following non-target strand cleavage (Cofsky et al., 2020; Naqvi et al., 2022; Singh et al., 2023). Overall, our mechanistic studies suggest a model in which Cas12a binding to seed mismatched targets is inhibited by Mg^2+^ but can be more readily overcome at low Mg^2+^ concentration; by contrast, mismatches in the PAM-distal region impair cleavage following target binding at low Mg^2+^ concentration (Fig. 6).

**Figure 6:**
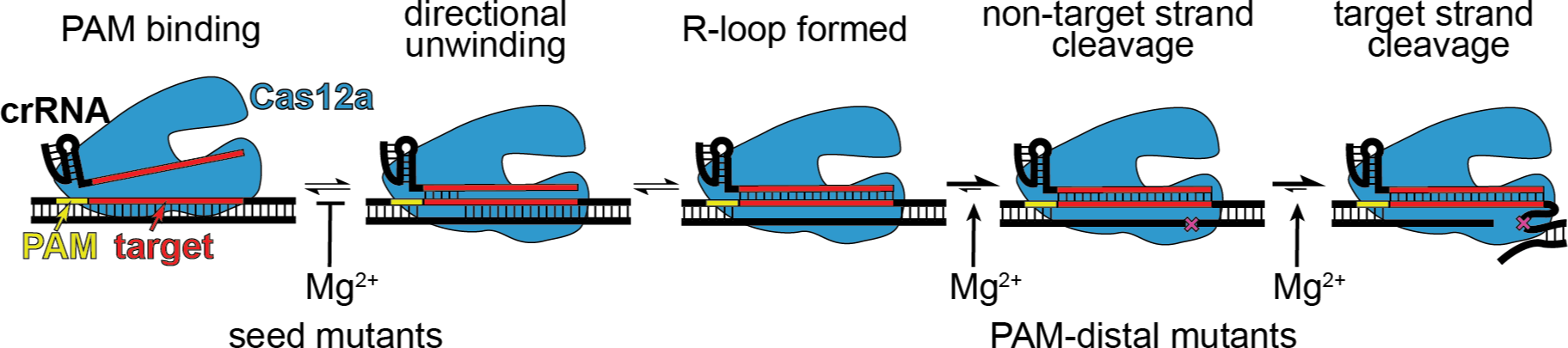
Model for Cas12a Mg^2+^-dependent steps with differential effects depending on mutant location. Mg^2+^ is inhibitory of target unwinding and binding when a seed mutation is present in the target. Mg^2+^-dependent conformational changes that occur prior to each cleavage step are likely impaired in the presence of PAM-distal mutations.

The metal-dependence of Cas effector specificity likely impacts the activity of these enzymes in native immune system settings. Metal ion availability may fluctuate during phage replication and transcription as nucleotides that coordinate Mg^2+^ are depleted. In our phage escape experiments, we observed that phages challenged with AsCas12a developed mutations throughout the PAM and seed. AsCas12a also had the lowest degree of specificity switching for seed mutants due to Mg^2+^ concentration. These results suggest that differences in Mg^2+^-dependent specificity may impact the types of mutants that emerge upon challenge by different Cas effectors. We also previously found that phage target sequences are prone to mutagenesis in the PAM-distal region when targeted by Cas12a (Schelling et al., 2023), potentially due to recombination-mediated repair following Cas12a target cleavage (Hossain et al., 2021; Wu et al., 2021). We now observe that single point mutations in the PAM-distal region are sufficient to allow escape from Fn and AsCas12a, potentially due to the strongly deleterious effect that these mutations are likely to have on second-strand cleavage under physiological metal ion conditions.

Our results suggest that specificity profiling performed at higher Mg^2+^ concentration may not reveal the true off-target sites for Cas effectors used in genome editing studies. Importantly, most *in vitro* genome-wide off-target site detection protocols suggest using a cleavage buffer containing 10 mM Mg^2+^ concentration (Cameron et al., 2017; Kim et al., 2016; Kleinstiver et al., 2016; Lazzarotto et al., 2020). Off-target sites that may be cleaved at lower Mg^2+^ concentration could be missed if specificity is only determined at a higher, non-physiological divalent metal ion condition. Conversely, use of higher Mg^2+^ concentration could lead to many false positive off-target results due to more rapid cleavage of some mismatched targets under these conditions. Thus, determination of off-target cleavage by Cas effectors at a range of metal ion conditions will be a critical addition to these well-established workflows.

## Data Availability

All scripts used for analysis of sequence data are available at github.com/sashital/Cas12a_metal_dependence. The fastq files generated from MiSeq sequencing for pLibrary cleavage products and escaped phage populations are available upon request and will be submitted to a repository.

## Acknowledgements

We thank members of the Sashital lab for helpful discussions and Michael Baker and David Wright from the Iowa State University DNA Facility for technical assistance for MiSeq data collection. This work was supported by NSF grant 1652661 and NIH grant GM140876 (D.G.S.).

## Methods

### Cas12a and crRNA purification

All plasmids, primers, and oligonucleotides are listed in Supplementary Data 2. Cas12a expression plasmids contained an N-terminal maltose binding protein tag containing a hexa-histidine affinity tag and tobacco etch virus (TEV) protease site followed by a codon optimized sequence for expression of Cas12a. All constructs were under control of a T7 RNA polymerase promoter and *lac* operator. The construct for LbCas12a expression (pMAL-His-LbCpf1-EC) - a gift from Jin-Soo Kim (RRID: Addgene 79008) (Kim et al., 2016) - was modified to delete sequence encoding an N-terminal nuclear localization sequence and to add a stop codon at the end of the LbCas12a sequence. All Cas12a proteins were expressed and purified using our previously described protocol for FnCas12a purification (Schelling et al., 2023).

*In vitro* transcription reactions to produce crRNAs targeting gene L and gene W targets were performed and RNA was purified as previously described (Schelling et al., 2023). Templates for this transcription reactions are listed in Supplementary Data 2. The crRNAs used for the ssDNA trans cleavage assay in Fig. S2C were chemically synthesized by Integrated DNA Technologies and are listed in Supplementary Data 2.

### Plasmid library preparation

DNA oligonucleotide pools were synthesized by Twist Bioscience containing target sequences with single nucleotide deletions, one mutation, or two mutations along with flanking regions on either end. The base sequences for the gene L and gene W oligonucleotide pools are provided in Supplementary Data 2. The oligonucleotide pools were amplified using Q5 DNA polymerase (New England Biolabs) in a 12 cycle PCR reaction. pUC19 plasmid was PCR amplified using primers containing homology to the oligo pools (Supplementary Data 2). Both PCR products were purified using Promega Wizard PCR purification kit. The PCR products were assembled using 75 ng of the backbone PCR DNA and 12.5 ng of the PCR amplified library in a 15 μL Gibson assembly reaction using NEBuilder HiFi DNA Assembly Master Mix (New England Biolabs) at 50 °C for 1 hr. The assembly reaction was transformed into NEB 5-alpha competent cells (New England Biolabs). After the recovery step, the entire recovery culture was used to inoculate 500 mL LB broth supplemented with 100 mg/mL ampicillin and incubated at 37 °C overnight with shaking. The overnight culture was cooled on ice for 30 min before harvesting. The plasmid library was isolated using QIAGEN Plasmid Midi Kit, aliquoted, and stored at -20 °C.

### Addition of spike-in plasmids to plasmid library

Our results suggest that mutant sequences in the plasmid library that cause significant cleavage defects may be non-specifically nicked by Fn or LbCas12a that has been activated by other sequences in the library (Fig. S2B). Thus, non-specific nicking may artifactually remove sequences from the uncleaved pool at higher magnesium ion conditions. To address this, we spiked in five additional plasmids to the plasmid library. These plasmids contain the same backbone as the library, but contain “target” sequences that bear no complementarity to the crRNAs and therefore do not undergo cleavage by Cas12a programmed with either gene L or gene W crRNA (Supplementary Data 2). Because the sequences are not specifically cleaved by Cas12a, they can be used as a reference to normalize sequences in the uncleaved fraction (see below). Non-specific cleavage by Fn and LbCas12a affects all sequences in the plasmid library, including the spike-in plasmids. Thus, normalization to the spike-in plasmids ensure that only specific cleavage by Cas12a is measured in our assay. Mutant sequences that are uncleaved by Cas12a should have similar abundance to the spike-in sequences.

For the spike-in sequences, five oligonucleotides containing the same flanking sequence as the libraries but with a random 24-nt sequence replacing the gene L or W target sequence were designed (Integrated DNA Technologies, Supplementary Data 2). These oligonucleotides were pooled, and a plasmid library was created as described above for the gene L and gene W oligonucleotide pools. The resulting library of random sequences was spiked into each plasmid library in a 1:100 ratio to create pLibrary for gene L or gene W.

### In vitro plasmid library cleavage assays and MiSeq sample preparation

Triplicate cleavage assays were prepared in reaction buffer (20 mM HEPES, pH 7.5, 100 mM KCl, 1 mM DTT, 5% glycerol and varying concentration of MgCl_2_). Cas12a RNP complex was formed by incubating Cas12a and crRNA at a 1:1.5 ratio at 37 °C for 10 min. To initiate cleavage, pLibrary (final concentration 15 ng/µL, preheated at 37 °C for 10 min) was added to Cas12a RNP (final concentration 50 nM). The reaction was incubated at 37 °C. 10 μL aliquots were quenched at 1 and 30 min by adding 10 µL 25:24:1 phenol-chloroform-isoamyl alcohol (Invitrogen). Two controls were included in which no protein or crRNA was added to the reaction or only Cas12a and no crRNA was added to the reaction. These controls contained reaction buffer with 10 mM MgCl_2_ and were incubated at 37 °C for the 30 min before adding phenol-chloroform-isoamyl alcohol. After extraction of the aqueous layer, DNA products were separated by electrophoresis on a 1% agarose gel. The gel was then stained with SYBR Safe (Invitrogen) and visualized using a UV transilluminator. The bands from the nicked and supercoiled fractions were excised and gel purified using QIAquick Gel Extraction Kit (Qiagen). Nextera adapters were added by performing a 25-cycle PCR reaction using primers bearing the Nextera adapters on the 5′ ends (Supplementary Data 2). These PCR products were cleaned up using QIAquick PCR Purification Kit (Qiagen) and used as a template for an 8-cycle PCR reaction to add barcodes for sample identification. Q5 DNA polymerase (New England Biolabs) was used for all PCR reactions. Samples were pooled and gel purified using the Promega Wizard PCR purification kit. Gel purified samples were analyzed on an Agilent 2100 Bioanalyzer and then submitted for Illumina MiSeq high-throughput sequencing at the Iowa State DNA Facility.

To quantify the fraction uncleaved and fraction nicked from agarose gel images, the intensity of the negatively supercoiled, nicked and linear bands was quantified using densitometry in ImageJ (Schneider et al., 2012). The fraction uncleaved and nicked were determined using the following equations:

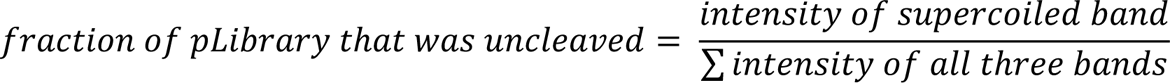

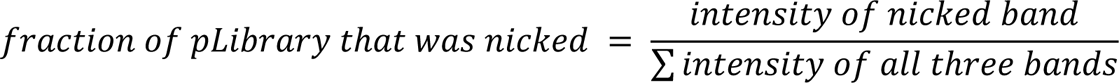

The fraction nicked values from the three replicates were averaged and used as a normalization factor (NF) for sequences in the nicked fraction (see below).

### Analysis of plasmid library cleavage

Python scripts used for analysis of plasmid library sequencing data are available at github.com/sashital/Cas12a_metal_dependence. The scripts were written in collaboration with ChatGPT (OpenAI, April-December, 2023 models) and were extensively validated.

For all data sets, R1 and R2 reads were joined using ea-utils (Aronesty, 2011, 2013). Only reads with identical sequences in the overlapping region of R1 and R2 were considered in our analysis.

Sequencing fastq files were first analyzed to determine the number of reads for each sequence present in the given pLibrary. The number of reads for each sequence in a given sample was normalized to the total number of reads in that sample to yield fraction of total reads (FTR):

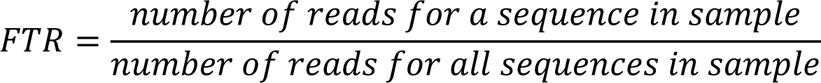

For each experimental file, the FTR for a given sequence was normalized to the FTR of the same sequence in the control files to yield normalized to control (NTC) values.

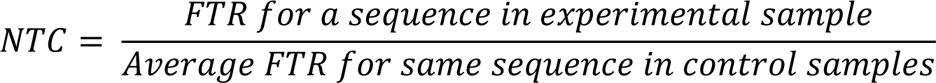

This normalization ensures that the original abundance of sequences in the library is accounted for when considering their abundance in the experimental samples.

For the uncleaved fraction (negatively supercoiled), the NTC values were further normalized to the spike-in sequences to determine the fraction uncleaved (*f*_uncleaved_) for each sequence:

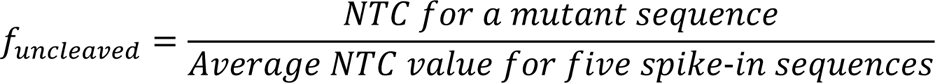

As described above, this normalization ensures that any non-specific cleavage that may have occurred for random sequences is accounted for when determining sequence abundance in the uncleaved fraction.

For the nicked fraction, the NTC values were further normalized using the normalization factor (NF) calculated from the average fraction of pLibrary that was nicked (described above). The following equation was used to determine the fraction nicked (*f*_nicked_) for each sequence:

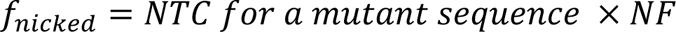

This normalization allows more direct comparison between conditions and orthologs by ensuring that samples that contained more nicked DNA have higher abundance of nicked sequences. This information is otherwise lost due to the PCR amplification of the nicked fractions.

For volcano plots, log2 fold-change for the fraction uncleaved or nicked was calculated as follows:

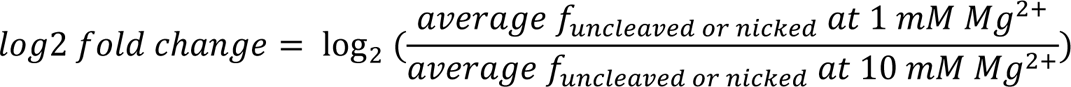

*P* values for fraction uncleaved or nicked values at 1 and 10 mM Mg^2+^ were determined using an unpaired, two-tailed t-test, and the log of these values were plotted on the volcano plots.

The fraction uncleaved and fraction nicked values were used to generate the heatmaps shown in Figure 2 and Supplementary Data 1. The fraction of DNA that was linearized was inferred as the fraction of DNA that was neither uncleaved nor nicked. Thus, the fraction of each sequence that was linearized (*f*_linear_) was calculated using:

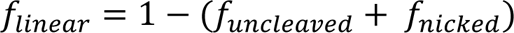

For each mutant sequence, the *f*_linear_ values versus Mg^2+^ concentration were fit to a linear regression to determine a slope. These values were used to generate the slope heatmaps.

### Cas12a collateral cleavage assays

Cleavage assays were prepared in reaction buffer (20 mM HEPES, pH 7.5, 100 mM KCl, 1 mM DTT, 5% glycerol and varying concentration of MgCl_2_). The Cas12a-crRNA RNP was first formed by incubating at 37 °C for 10 min at a final concentration of 20 nM Cas12a and 30 nM crRNA. For cleavage assays in which two plasmids were mixed, the empty pUC19 was added to a final concentration of 15 ng/μL and the activator target or plasmid library was added to a final concentration of 1.5 ng/μL. The DNA was preheated at 37 °C for 10 min and then mixed with RNP to initiate cleavage. Aliquots were quenched at the indicated time points by adding an equal volume of phenol-CHCl_3_-isoamyl alcohol and analyzed by agarose gel electrophoresis as described above.

The ssDNA collateral cleavage assays were performed similarly, but with the following modifications. The Cas12a-crRNA RNP was incubated with a short dsDNA activator (PS4 sequence from (Murugan et al., 2020), final concentration 30 nM) at 37 °C for 10 min before being added to M13mp18 single-stranded DNA (New England Biolabs, final concentration 25 ng/μL). Aliquots were quenched at the indicated time points by adding an equal volume of phenol-CHCl_3_-isoamyl alcohol and analyzed by agarose gel electrophoresis as described above.

### Time-course cleavage assays of individual target sequences

All cleavage assays summarized in Figure 4 and S4 were performed in reaction buffer (20 mM HEPES, pH 7.5, 100 mM KCl, 1 mM DTT, 5% glycerol) to which the indicated concentration of MgCl_2_ was added at varying times as described below. Cas12a RNP complex was formed by incubating Cas12a and crRNA at a 1:1.5 ratio (final concentration of 50 nM:75 nM or 100 nM:150 nM) at 37 °C for 10 min.

For reactions that were initiated by adding DNA, both the RNP and DNA target plasmid were diluted in reaction buffer containing the indicated concentration of MgCl_2_. The DNA plasmid was preheated for 10 min at the appropriate temperature and then added at a final concentration of 15 ng/μL to Cas12a RNP. Aliquots were quenched at the indicated time points using phenol-CHCl_3_-isoamylalcohol and analyzed by agarose gel electrophoresis as described above.

For reactions that were initiated by adding MgCl_2_, both RNP and DNA target plasmid were diluted in reaction buffer without MgCl_2_. After pre-incubating both RNP and DNA individually at 37 °C for 10 min, the RNP and DNA were mixed to the final concentrations of 50 nM RNP and 15 ng/μL DNA and incubated at 37 °C for 30 min to allow binding (final volume 80 μL). 20 μL of a 1X reaction buffer containing 50 mM or 5 mM MgCl_2_ was then added to initiate cleavage, resulting in a final MgCl_2_ of 10 mM or 1 mM MgCl_2_. Aliquots were quenched at the indicated time points using phenol-CHCl_3_-isoamylalcohol and analyzed by agarose gel electrophoresis as described above.

For binding time course reactions in which Cas12a RNP was allowed to bind to DNA in the absence of MgCl_2_ before initiating cleavage by mixing with MgCl_2_, both RNP and DNA target plasmid were diluted in reaction buffer without MgCl_2_ and incubated individually at 37 °C for 10 min. The RNP and DNA were mixed to the final concentrations of 50 nM RNP and 15 ng/μL DNA at 37 °C. Following mixing, 10 μL aliquots were taken at 7 s, 15 s, 30 s, 1 min, 2 min, 5 min, 15 min and 30 min. The aliquots were initially mixed with 10 μL of 1X reaction buffer supplemented with 20 mM MgCl_2_ at 37 °C for 5 s to allow any DNA bound by RNP to be cleaved. Cleavage was quenched immediately after by mixing 10 μL of the reaction with 10 μL phenol-CHCl_3_-isoamylalcohol. The samples were analyzed by agarose gel electrophoresis as described above.

The gel images were quantified by densitometry using ImageJ to determine the intensity of the negatively supercoiled, nicked and linear bands. To differentiate first-versus second-strand cleavage, we reasoned that all nicked DNA (i.e. DNA that has undergone the first cleavage event) eventually becomes linearized. Thus, first-strand cleavage was determined by adding together the nicked and linear bands:

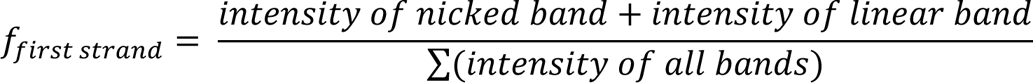

where *f_first strand_* is the fraction of DNA that has undergone first-strand cleavage. DNA that has undergone second-strand should be linearized, and was determined using:

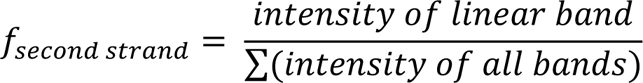

where *f_second strand_* is the fraction of DNA that has undergone second-strand cleavage. These values were plotted versus time and fit to a single- or double-exponential rate equation in GraphPad Prism. All experiments were performed at least three times, and rate curves show the average of three replicates with error bars representing standard deviation.

Rate constants reported in Figure 1C and normalized rate constants reported in Figure 3D were derived from fits of individual replicates to a single-exponential rate equation in GraphPad Prism. For normalized rate constants, the *k*_obs_ values for three replicates at each Cas12a concentration were averaged, and then each average value was divided by the average *k*_obs_ value at 50 nM Cas12a. Standard deviation values for *k*_obs_ was propagated to determine error for these normalized *k*_obs_ values.

### Phage competition and escape assays

*E. coli* strain BW25113 (Baba et al., 2006) and a virulent strain of lambda phage (λ_vir_) (Jacob and Wollman, 1954) or mutants of these strains (Schelling et al., 2023) were used for all experiments involving phage infection. Plasmids for expression of Cas12a orthologs and crRNAs are listed in Supplementary Data 2. For competition assays, FnCas12a was expressed from a arabinose-inducible pBAD promoter using a previously described modified pACYC vector (Schelling et al., 2023). For phage escape assays, the Cas12a orthologs were expressed from a weak constitutive promoter using a modified pACYC vector that we have previously shown provides decreased Cas12a defense (Schelling et al., 2023), allowing for more rapid phage escape. All crRNAs were expressed from CRISPR constructs under control of an arabinose-inducible pBAD promoter in a modified pUC19 vector.

For all phage assays, overnight cultures were started using a single colony of *E. coli* BW25113 transformed with Cas12a and crRNA expression plasmids in LB media with ampicillin and chloramphenicol added for plasmid selection. The next day, these overnight cultures were used to inoculate cultures at a 1:100 dilution in LB media containing 100 µg/mL ampicillin, 25 µg/mL chloramphenicol, 20 mM arabinose, and 10 mM MgSO_4_ or 150 mM MgSO_4_ for data shown in Fig. 5G. Cultures were grown in a TECAN infinite M Nano+ 96 well plate reader at 280 rpm and 37 °C and OD measurements at 600 nm wavelength were measured every 10 minutes. Phage was added to each well at OD_600_ 0.4 at the specified MOI.

For competition assays the A2T and G17T gene L mutant lambda phages (Schelling et al., 2023) were mixed at the indicated ratio and diluted by 10^6^ (Fig. 1B) or 10^7^ (Fig. 5G) via serial dilution in LB media. These dilute mixtures (3 μL) were used to infect 150 μL cultures at OD_600_ of 0.4.

For phage escape experiments in which time points were taken, 20 µL samples were taken from each well (initially containing 200 μL) at various time points and stored at 4 °C. Reported OD measurements were corrected to compensate for the reduction in volume.

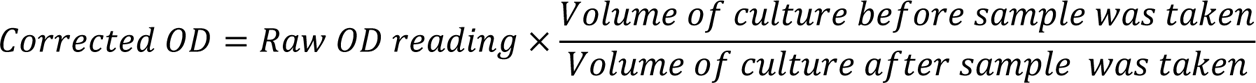

Once all samples were collected, the samples were centrifuged at 15,000 RPM for 5 minutes and 5 µL of supernatant was used as a template for adapter and barcode PCRs as previously described (Schelling et al., 2023). The samples were pooled and sequenced using MiSeq.

### Analysis of mutant phage populations

For competition assays, R1 and R2 reads were joined using ea-utils (Aronesty, 2011, 2013). Only reads with identical sequences in the overlapping region of R1 and R2 were considered in our analysis. The number of reads containing the A2T or G17T mutation was determined by searching for the mutant sequence using grep -c. These values were divided by the total number of reads in the sample to determine the fraction of reads containing each mutation. Statistical significance for the 24 replicates performed at 10 and 150 mM MgSO_4_ was tested in GraphPad Prism using an unpaired, nonparametric Komogorov-Smirnov test to compare the cumulative distributions of the G17T mutant in each sample.

Python scripts used to analyze mutant phage populations in phage escape assays are available at github.com/sashital/Cas12a_metal_dependence. Percent mutated, nucleotide diversity, and Z-scores for three experimental replicates were calculated as previously described (Schelling et al., 2023), with the following adjustment. To calculate nucleotide diversity, we considered all mutated sequences present across three replicates for a given Cas12a ortholog/crRNA pair, rather than calculating a diversity score for each individual replicate. This strategy better represents the potential diversity that can arise through random mutant emergence that may occur in different cultures. Deletions were accounted for and treated as mismatches relative to the perfect target, and deletions of multiple nucleotides were treated as multiple mismatches. Average nucleotide diversity scores were determined for each ortholog by averaging the nucleotide diversity score for every culture in which mutants arose (all eight genes for AsCas12a and FnCas12a, five of eight genes for LbCas12a). The error is reported as standard error of the mean, and significance was tested using an unpaired two-tailed *t* test.

### Western blot detection of Cas12a expression levels

A single colony of *E. coli* BW25113 with Cas12a and gene W crRNA expression plasmids was grown overnight in LB media supplemented with 100 μg/mL ampicillin, and 25 μg/mL chloramphenicol. LB supplemented with same amount of antibiotics, 20 mM arabinose, and 10 mM MgSO_4_ was inoculated with the overnight culture in a 1:100 ratio. The cultures were incubated at 37 °C with shaking. 1 mL samples of the cultures were harvested by centrifugation after 2 and 4 h of expression. The cell pellets were resuspended with water and then lysed with 2X SDS dye following by heating for 5 min at 95 °C. 10 μL of lysed cell samples and 10 μL of 100 nM purified Cas12a proteins were separated by SDS-polyacrylamide gel electrophoresis and transferred onto a nitrocellulose membrane (Cytiva). The membranes were blocked 1 h with 0.1% BSA in TBST buffer (20 mM Tris-HCl, 150 mM NaCl pH 7.6 and 0.05% Tween 20). After washing three times with TBST, the membranes were incubated with the relevant primary antibody anti-FnCpf1 (GenScript), anti-LbCpf1 (Millipore), or anti-AsCpf1 (Millipore) in 10 mL of BSA in TBST for 1 h at room temperature. A second blot containing the same sample was incubated with anti-GAPDH (Invitrogen) to serve as a loading control for the lysates. The membranes were washed three time with TBST buffer before incubating with goat anti-mouse secondary antibody (Fisher) in 10 mL of BSA in TBST for 1 h at room temperature. Final washing was conducted three time before imaging the blot with a chemiluminescent substrate (Pierce).

## Figures and Legends

**Figure S1:**
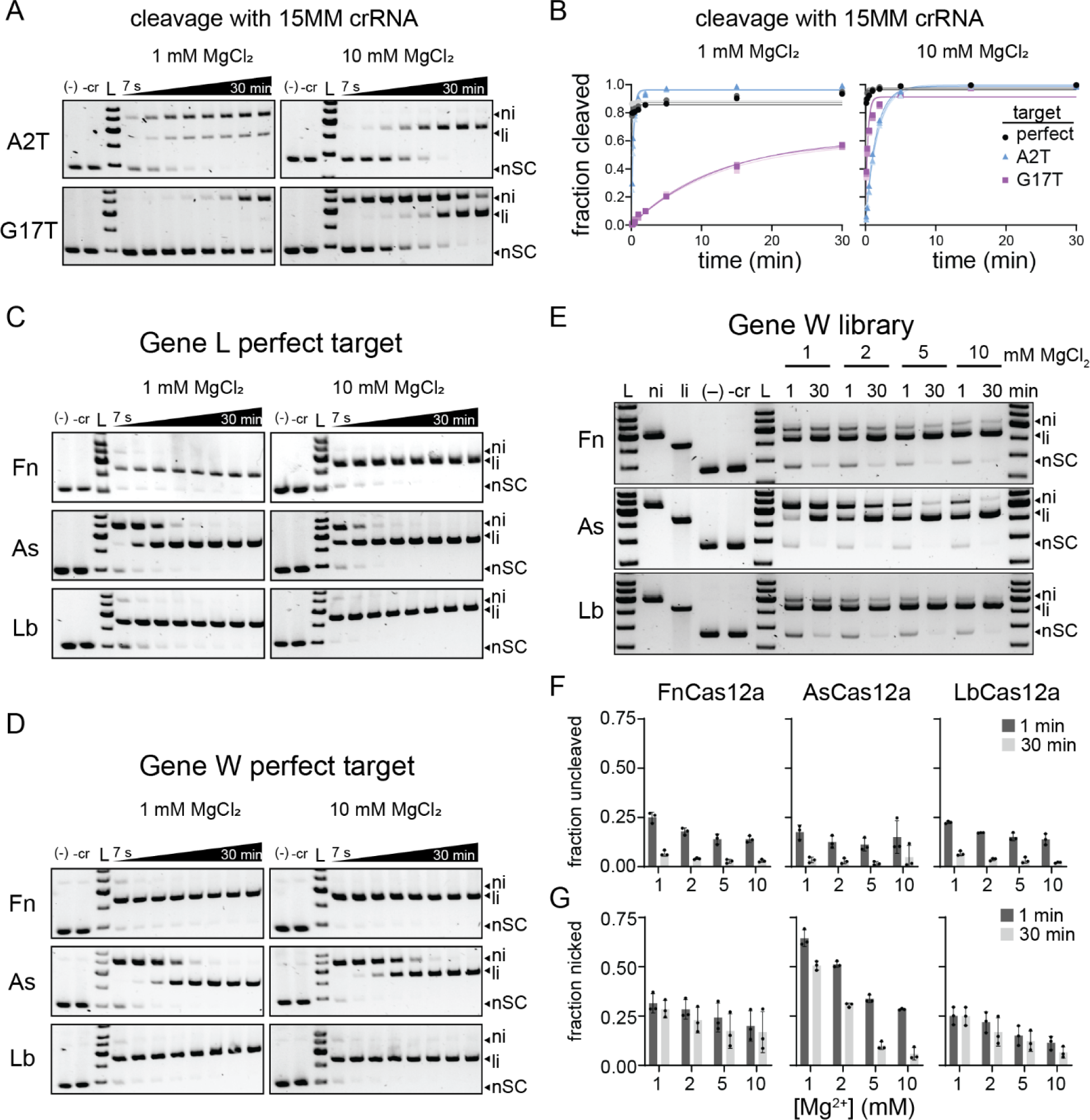
Cleavage of gene L and W targets and gene W target library (Related to Figure 1) A-B) Cleavage and quantification of mutant gene L targets by FnCas12a bearing a crRNA containing a mismatch at position 15. For agarose gels in (A), the time points are 7 s, 15 s, 30 s, 1 min, 2 min, 5 min, 15 min, and 30 min. ni = nicked, li = linear, nSC = negatively supercoiled. (–) lane contains no protein, -cr contains protein but no crRNA. Both controls contained the indicated MgCl_2_ concentration and were incubated at 37 °C for 30 min. Each gel is representative of three replicates. The triplicate gels are quantified in (B) and fit to a single-exponential rate equation to derive the rate constants reported in Fig. 1C. Cleavage of the perfect gene L target cleaved by FnCas12a bearing the mismatched crRNA is also shown for comparison. C-D) Cleavage of the perfectly matched gene L (C) and gene W (D) targets by each Cas12a ortholog at 1 and 10 mM MgCl_2_. Timepoints and labels are as in (A). Both controls contained 10 mM MgCl_2_ and were incubated at 37 °C for 30 min. Gels are representative of three replicates. For FnCas12a cleaving gene L, the same gels are shown in Fig. S5A. E) Gene W plasmid library cleavage at four Mg^2+^ concentrations for FnCas12a (Fn), AsCas12a (As), and LbCas12a (Lb). ni = nicked, li = linear, nSC = negatively supercoiled. (–) lane contains no protein, -cr contains protein but no crRNA. Both controls contained 10 mM MgCl_2_ and were incubated at 37 °C for 30 min. Gel is representative of three replicates. F-G) Quantification of fraction uncleaved (F) or nicked (G) for the gene W library. The average of three replicates is plotted, with individual data points shown as dots and error bars representing standard deviation.

**Figure S2:**
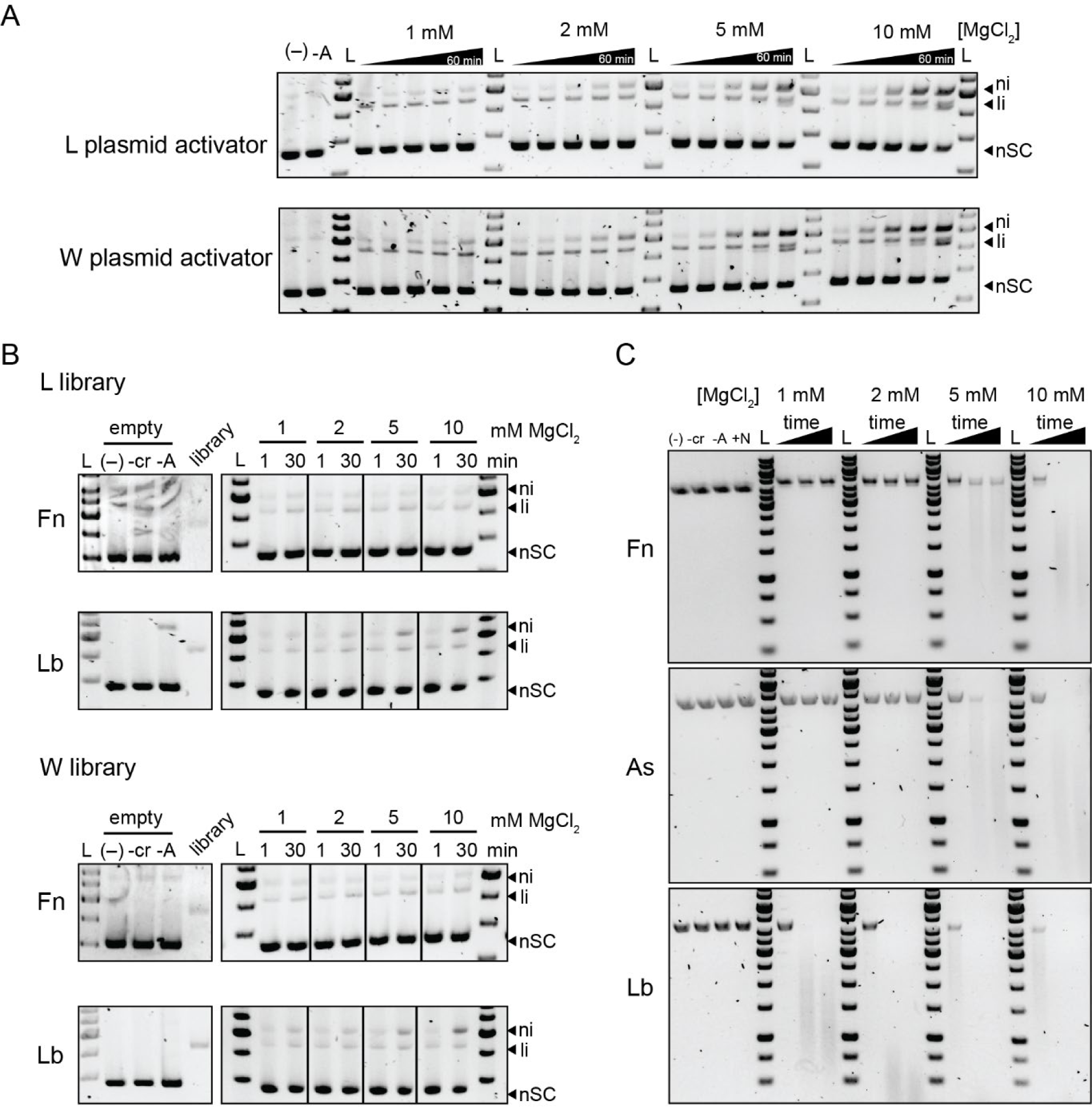
Collateral cleavage by Cas12a is Mg^2+^-dependent (Related to Figure 1) A) Cleavage of an empty pUC19 plasmid that lacks any complementarity to the crRNA by LbCas12a activated with either a gene L and gene W target pUC19 plasmid. The empty plasmid was in 10X excess of the activator target plasmid. The target plasmid was linearized by the first time point, and additional cleavage products observed at subsequent time points are most likely due to cleavage of the empty pUC19 plasmid. Time points: 1 min, 5 min, 15 min, 30 min, 60 min. The first two lanes are controls with empty pUC19 plasmid without any protein added (–) or with Cas12a-crRNA but without the activator target plasmid added (-A) at 10 mM MgCl_2_ incubated at 37 °C for the longest time point. B) Cleavage of an empty pUC19 plasmid that lacks any complementarity to the crRNA by Fn or LbCas12a activated with either the gene L and gene W target plasmid library. The empty plasmid was in 10X excess of the plasmid library. The gels on the left contain controls with empty pUC19 plasmid without any protein added (–), with Cas12a but no crRNA (-cr) or with Cas12a-crRNA but with only the empty pUC19 plasmid (-A) or the plasmid library (library) at 10 mM MgCl_2_ incubated at 37 °C for 30 min. C) Cleavage of single-stranded M13 phage DNA by Cas12a activated with a short double-stranded DNA activator. The time points were 15 s, 30 min, and 60 min. The first four lanes are controls with M13 DNA without any protein added (–), M13 DNA with only Cas12a added (-cr), Cas12a-crRNA but without the activator DNA added (-A), or Cas12a-crRNA with a non-targeting dsDNA oligo added (+N). All controls contained 10 mM MgCl_2_ and were incubated at 37 °C for the longest time point.

**Figure S3:**
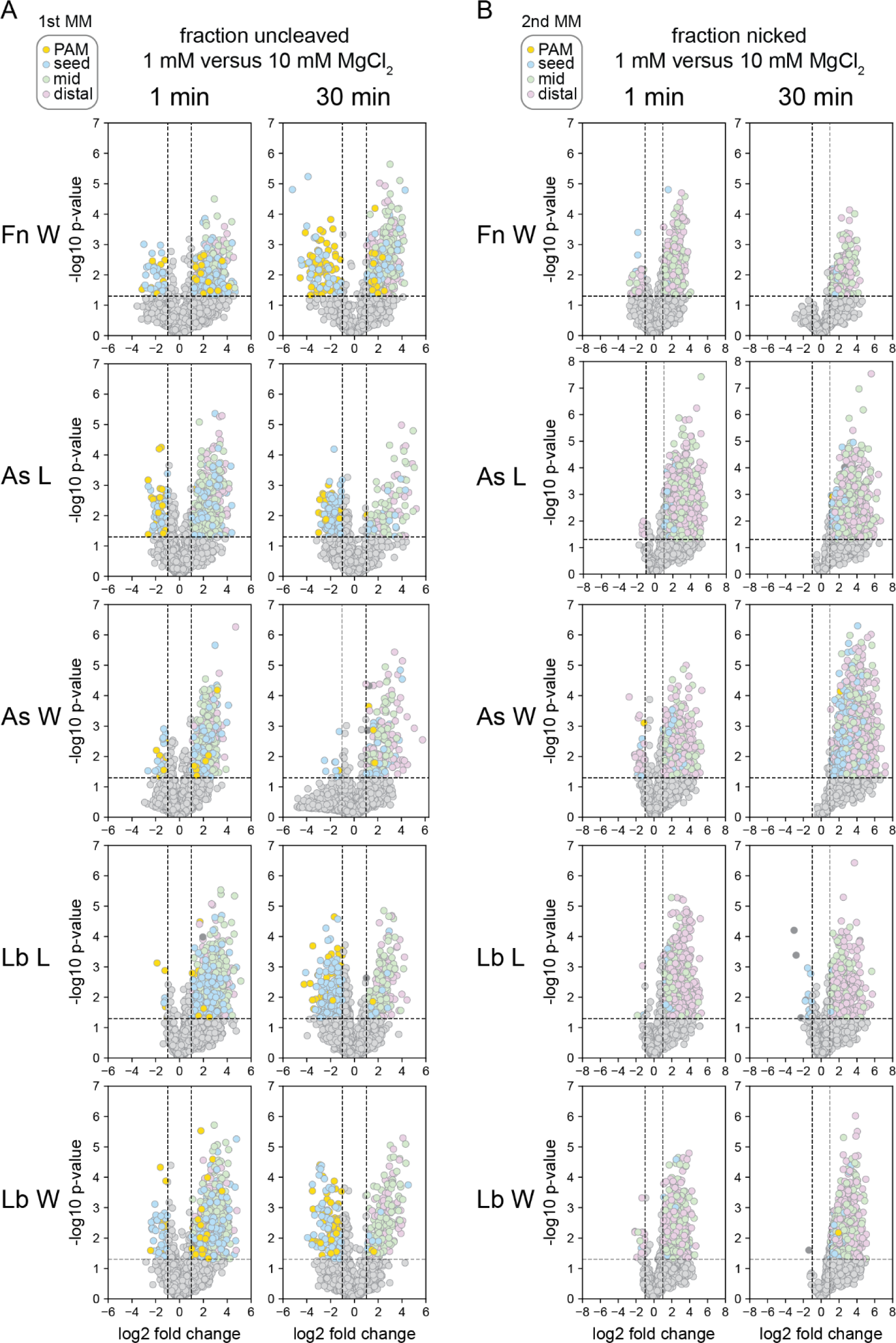
Volcano plots for uncleaved and nicked fractions for all Cas12a orthologs and libraries. (A-B) Volcano plots comparing sequences present in uncleaved (A) or nicked (B) fractions following cleavage at 1 or 10 mM MgCl_2_ for 1 or 30 min. Data points in (A) are colored by the location of the first mutation in the sequence. The plots for each ortholog FnCas12a (Fn), AsCas12a (As) or LbCas12a (Lb) and the gene L or W library are shown. Plots for gene L cleavage by FnCas12a are shown in Fig. 2A-B. Data points in (B) are colored by the location of the second mutation in the sequence. *P* values compare up to three replicates using an unpaired two-tailed *t* test

**Figure S4:**
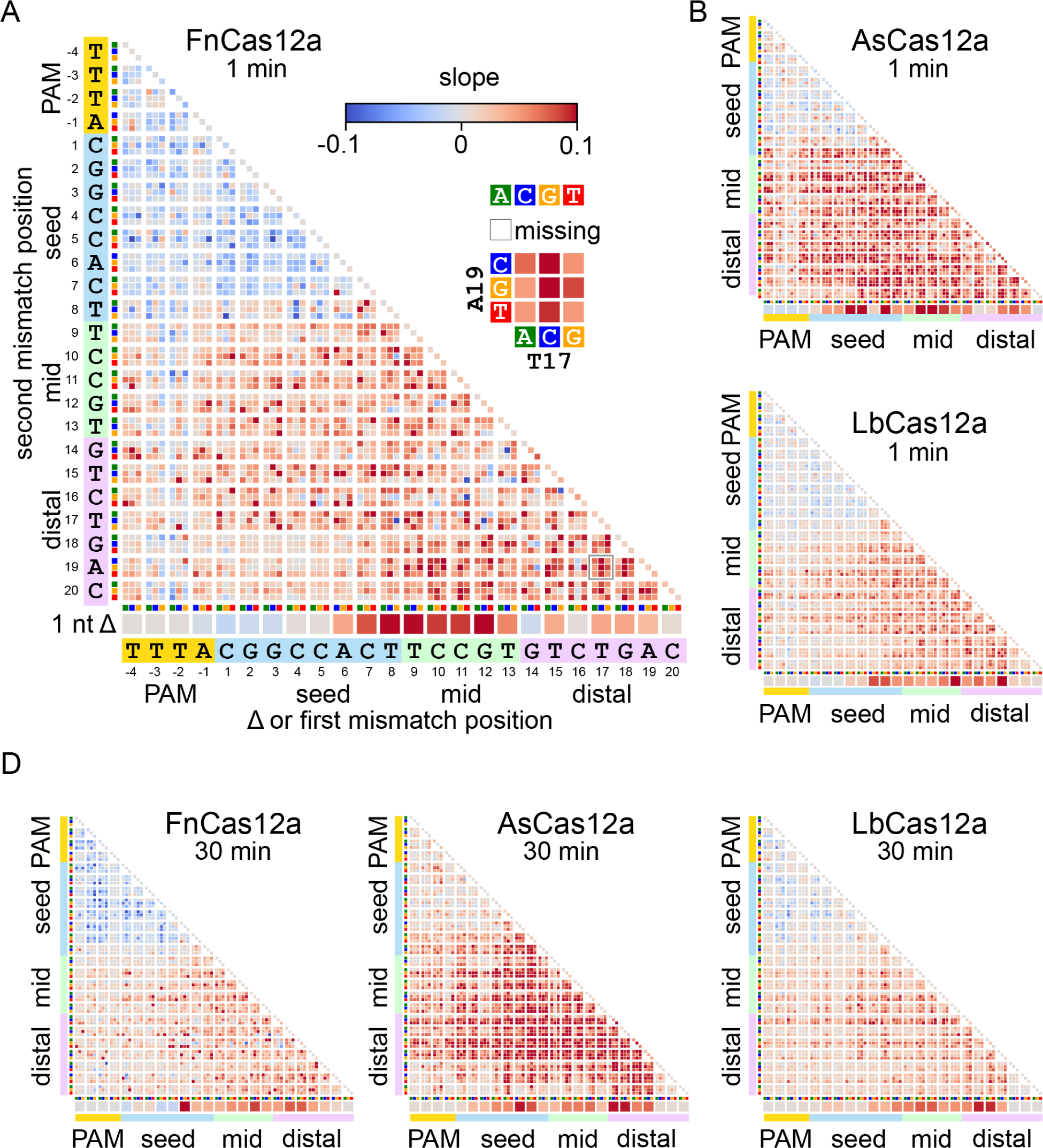
Slope heatmaps for the gene W target (Related to Figure 3) A) Heatmap plotting slopes for fully cleaved DNA versus Mg^2+^ as determined in Figure 3A-B. The heatmap is for the gene W target plasmid library following 1 min cleavage by FnCas12a. Missing sequences are represented by white boxes. B) Slope heatmaps of the gene W target plasmid library following 1 min cleavage by AsCas12a or LbCas12a. C) Slope heatmaps of the gene W target plasmid library following 30 min cleavage by all three orthologs.

**Figure S5:**
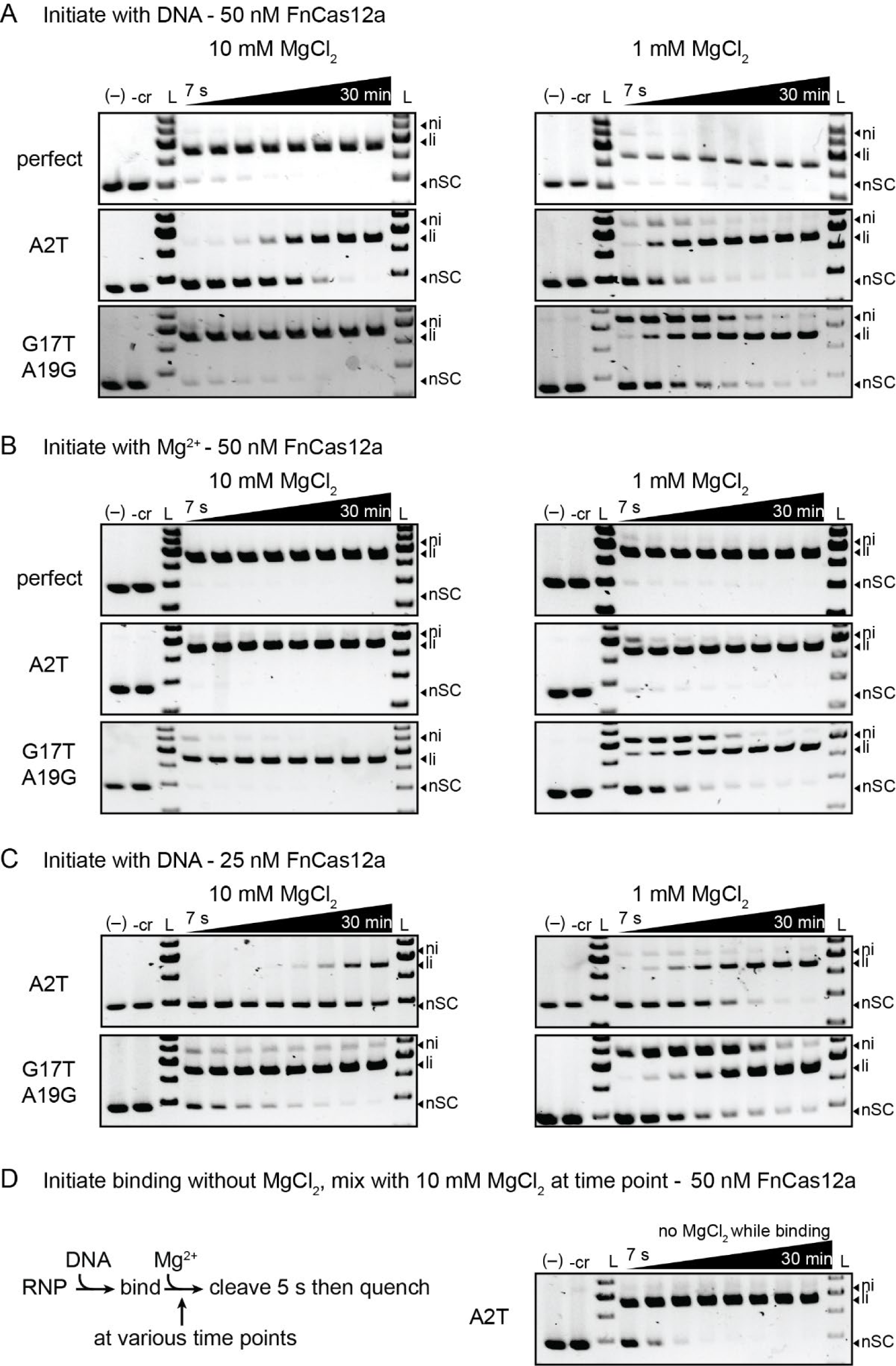
Representative agarose gels for cleavage assays quantified in Figure 4 (Related to Figure 4) Cleavage of plasmids bearing the indicated gene L target sequence by FnCas12a for the conditions described below. The gene L sequence either perfectly matched the crRNA (perfect) or contained a seed (A2T) or two PAM-distal (G17T A19G) mutations. For all gels, (–) indicates a control in which no protein was added to the DNA and -cr indicates a control in which only Cas12a without crRNA was added to DNA before incubating at 37 °C for 30 min. Controls were performed at the same MgCl_2_ concentration as the cleavage reaction. The time points for all gels were 7 s, 15 s, 30 s, 1 min, 2 min, 5 min, 15 min and 30 min. Each gel is a representative of at least three replicates. ni = nicked, li = linear, nSC = negatively supercoiled. A) Gels associated with the quantified data shown in Figure 4B and 4D (50 nM RNP) and 4E (A2T mutant) in which Cas12a cleavage was initiated by mixing Cas12a-crRNA RNP together with DNA in the presence of the indicated concentration of MgCl_2_. The gels for the perfect target are also shown in Fig. S1C. B) Gels associated with the quantified data shown in Figure 4C in which Cas12a-crRNA RNP was first incubated with DNA for 30 min prior to addition of the indicated concentration of MgCl_2_ to initiate cleavage. C) Gels associated with the quantified data shown in Figure 4D (25 nM RNP) in which Cas12a cleavage was initiated by mixing Cas12a-crRNA RNP together with DNA in the presence of the indicated concentration of MgCl_2_. D) Gels associated with the quantified data shown in Figure 4E in which binding was initiated in the absence of MgCl_2_. A schematic on the left describes how these reactions were performed. An RNP-DNA binding reaction was initiated in the absence of MgCl_2_. At each time point, an aliquot from the binding reaction was mixed with 10 mM MgCl_2_ for 5 s, followed by immediate quenching.

**Figure S6:**
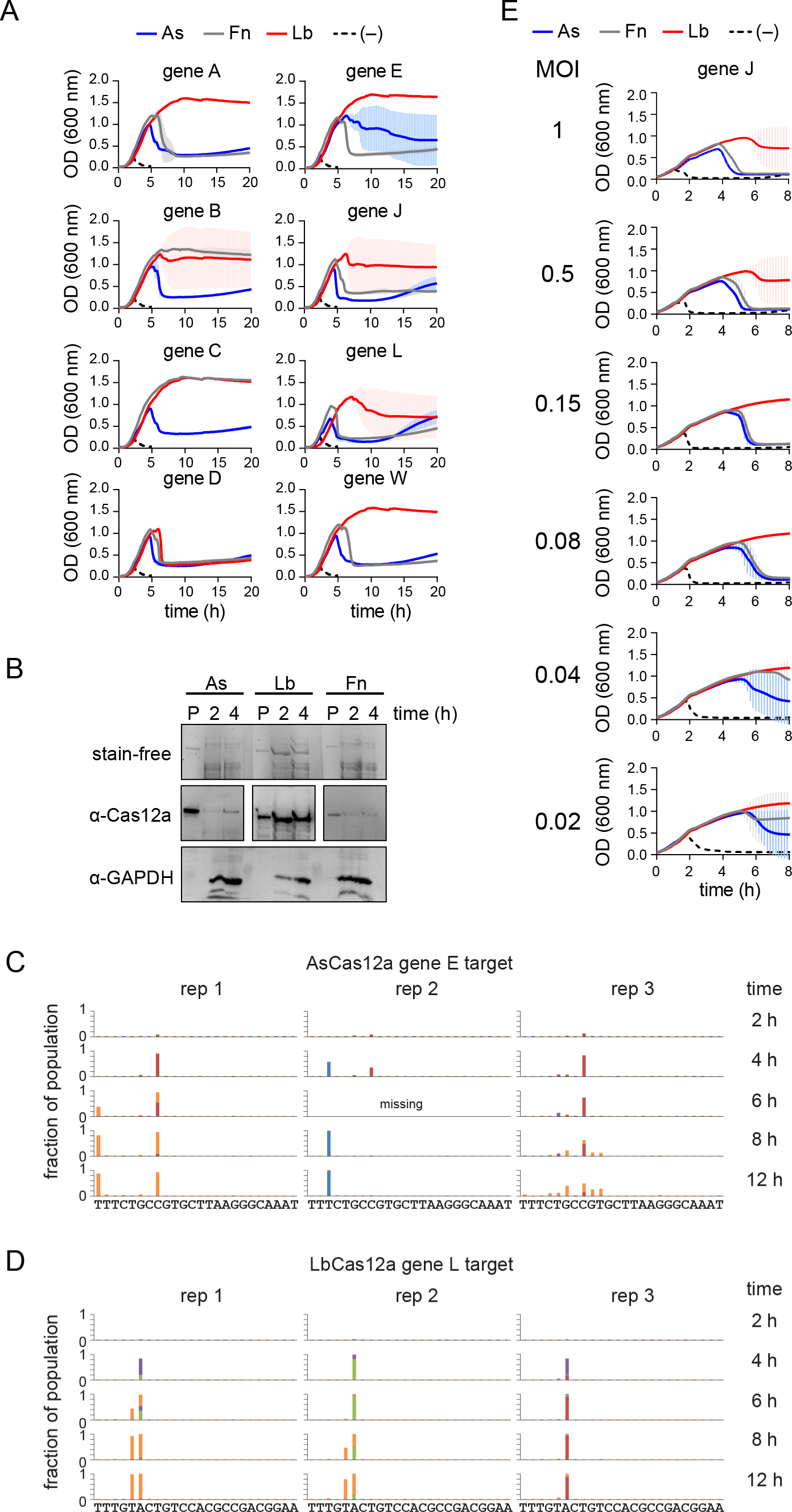
Growth curves, Western blots and individual replicates for phage escape experiments (Related to Figure 5) A) Growth curves (optical density at 600 nm wavelength vs. time) for *E. coli* cultures expressing each Cas12a ortholog and a crRNA targeting the indicated gene. Cultures were infected with phage λ_vir_ at an OD_600_ of 0.3. The average of three replicates are shown and error bars represent standard deviation. Large error bars indicate that some replicates did not undergo lysis while some did. B) Western blot analysis of Cas12a ortholog expression in *E. coli* cultures used for phage escape assays. The top panel shows the SDS-PAGE gel visualized using 2,2,2-trichloroethanol stain-free imaging. Purified AsCas12a, FnCas12a or LbCas12a (∼150 ng) was loaded in the first of three lanes, followed by lysates of cultures expressing each ortholog harvested at the indicated time points. The middle panel shows the Western blots using antibodies against each individual Cas12a ortholog. The bottom panel shows a Western blot using an antibody against *E. coli* GAPDH as a loading control for the lysates. C-D) Bar graphs showing double mutations that arose over time for three replicates of cultures expressing AsCas12a bearing a crRNA targeting gene E (C) or LbCas12a bearing a crRNA targeting gene L. A second mutation arose after an initial mutation for two of three replicates for each set of cultures. E) Growth curves for *E. coli* cultures expressing each Cas12a ortholog and a crRNA targeting gene J infected with the indicated MOI of phage λ_vir_ at an OD_600_ of 0.3. The average of three replicates are shown and error bars represent standard deviation. Large error bars indicate that some replicates did not undergo lysis while some did.

**Supplementary Data 1: Animated gifs showing the abundance of each mutant sequence in the uncleaved or nicked fraction.** Each gif shows all four Mg^2+^ concentrations for a given ortholog and gene at a given time point.

**Supplementary Data 2: All plasmids, primers and oligonucleotides used in this study.**

