## Supplementary Data 1 for "CRISPR-Cas12a exhibits metal-dependent specificity switching": README.txt

Description of animated gif files in this folder
As_L_1_uncleaved.gif: Heatmap plotting the fraction uncleaved values for the gene L library following 1 min cleavage by AsCas12a
As_L_30_uncleaved.gif: Heatmap plotting the fraction uncleaved values for the gene L library following 30 min cleavage by AsCas12a
As_W_1_uncleaved.gif: Heatmap plotting the fraction uncleaved values for the gene W library following 1 min cleavage by AsCas12a
As_W_30_uncleaved.gif: Heatmap plotting the fraction uncleaved values for the gene W library following 30 min cleavage by AsCas12a
Fn_L_1_uncleaved.gif: Heatmap plotting the fraction uncleaved values for the gene L library following 1 min cleavage by FnCas12a
Fn_L_30_uncleaved.gif: Heatmap plotting the fraction uncleaved values for the gene L library following 30 min cleavage by FnCas12a
Fn_W_1_uncleaved.gif: Heatmap plotting the fraction uncleaved values for the gene W library following 1 min cleavage by FnCas12a
Fn_W_30_uncleaved.gif: Heatmap plotting the fraction uncleaved values for the gene W library following 30 min cleavage by FnCas12a
Lb_L_1_uncleaved.gif: Heatmap plotting the fraction uncleaved values for the gene L library following 1 min cleavage by LbCas12a
Lb_L_30_uncleaved.gif: Heatmap plotting the fraction uncleaved values for the gene L library following 30 min cleavage by LbCas12a
Lb_W_1_uncleaved.gif: Heatmap plotting the fraction uncleaved values for the gene W library following 1 min cleavage by LbCas12a
Lb_W_30_uncleaved.gif: Heatmap plotting the fraction uncleaved values for the gene W library following 30 min cleavage by LbCas12a
