## Supplementary figures and images for "CRISPR-Cas12a exhibits metal-dependent specificity switching"

### As_L_1_nicked.gif

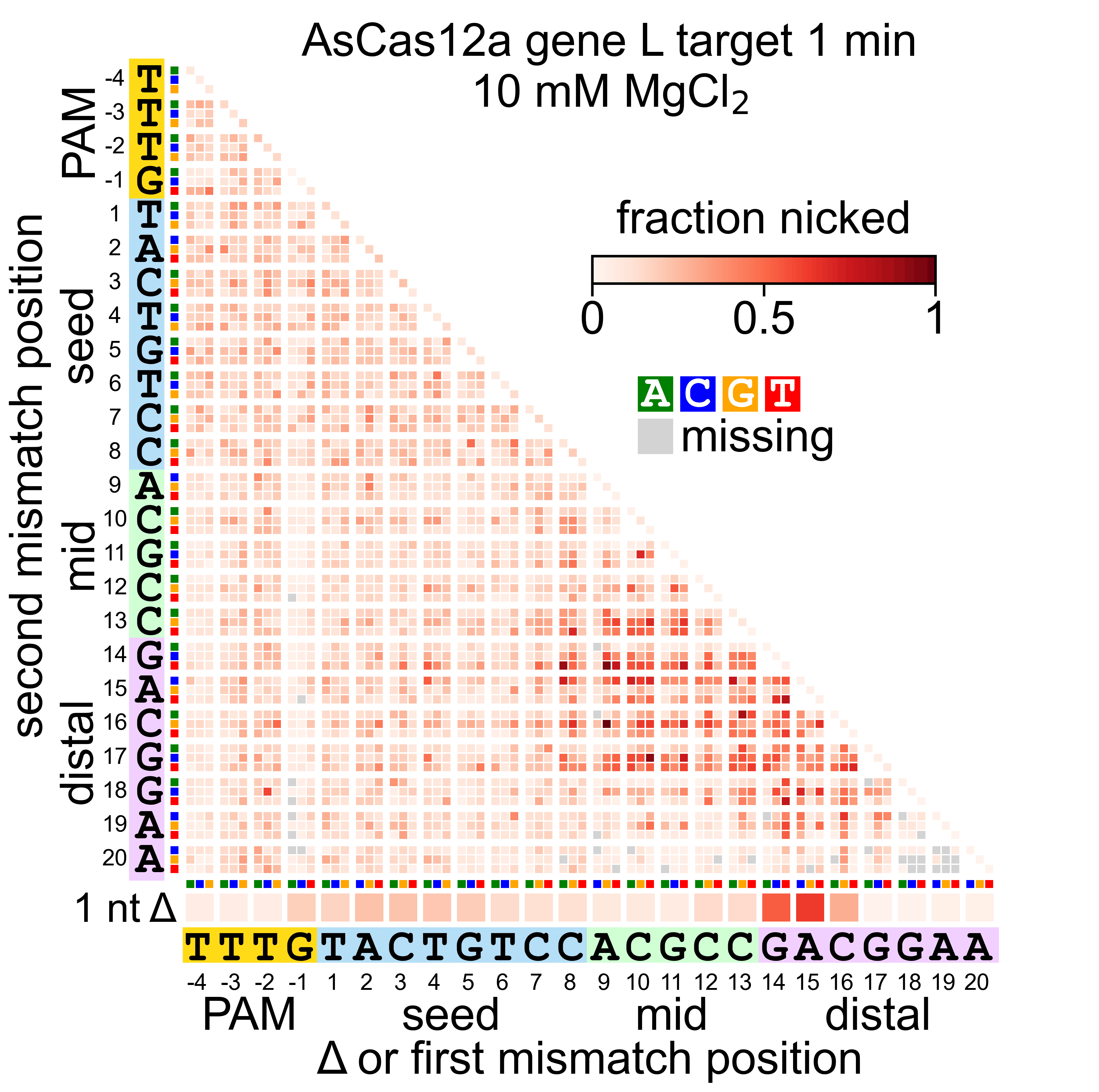

### As_L_1_uncleaved.gif

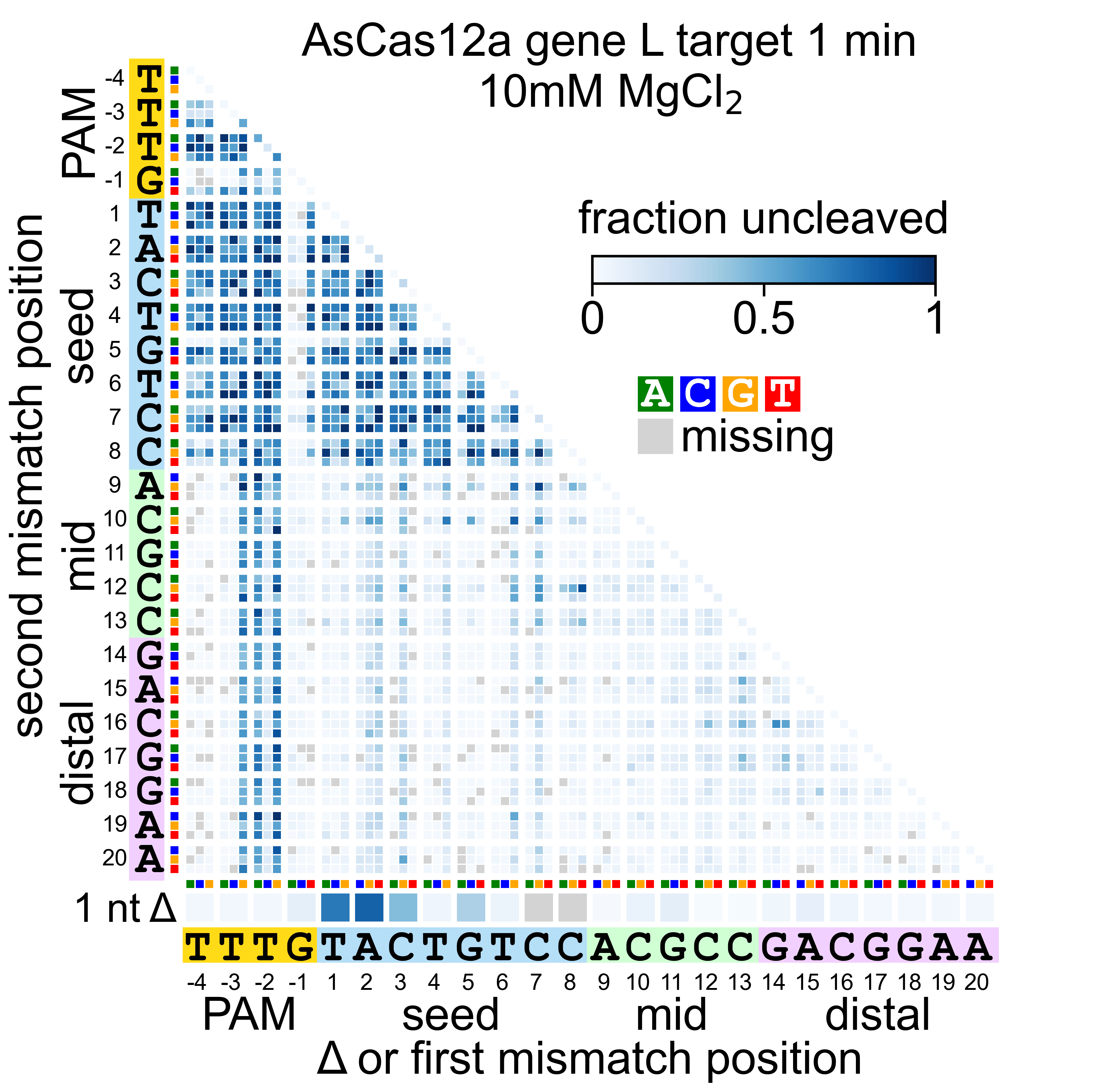

### As_L_30_nicked.gif

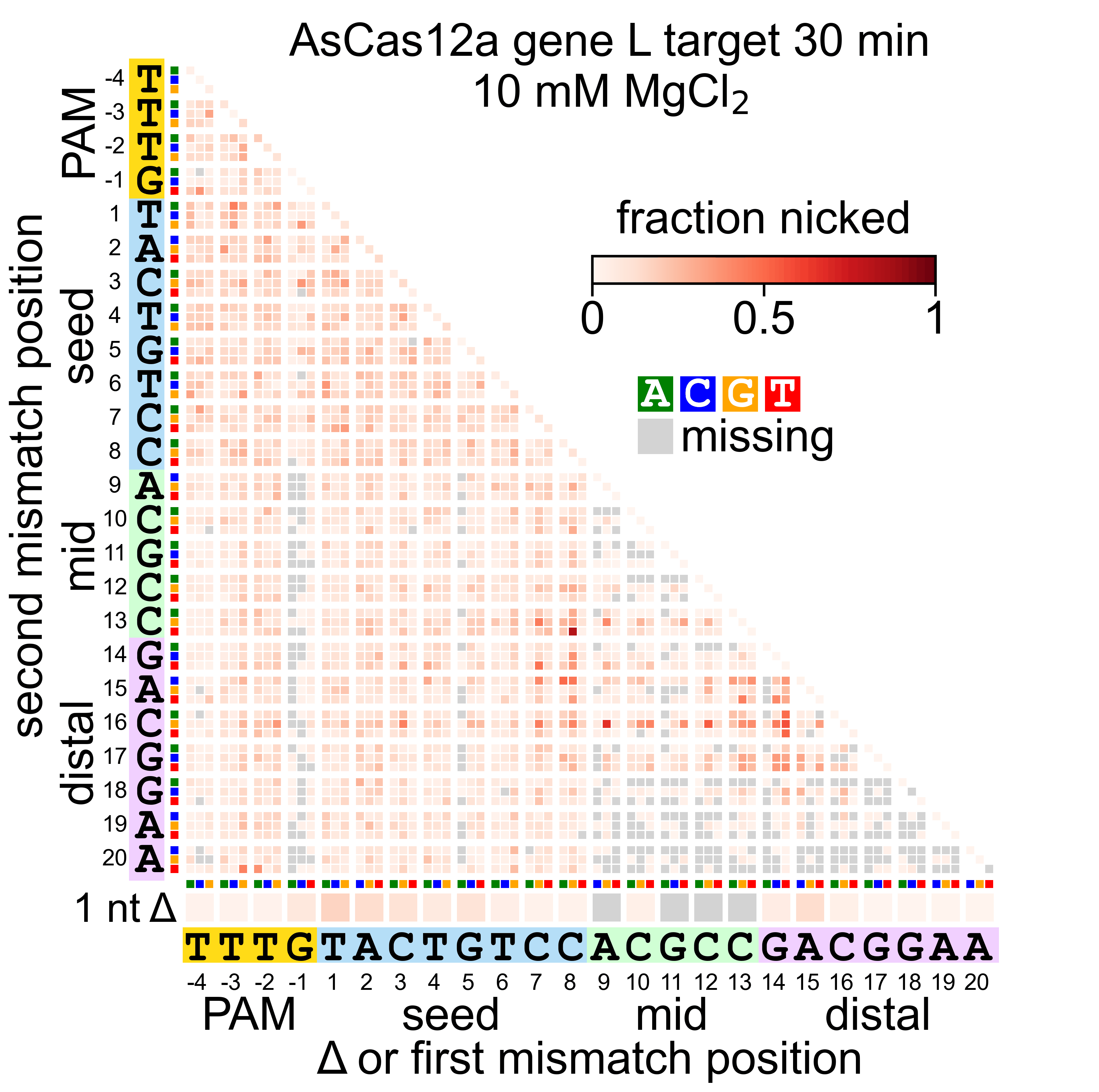

### As_L_30_uncleaved.gif

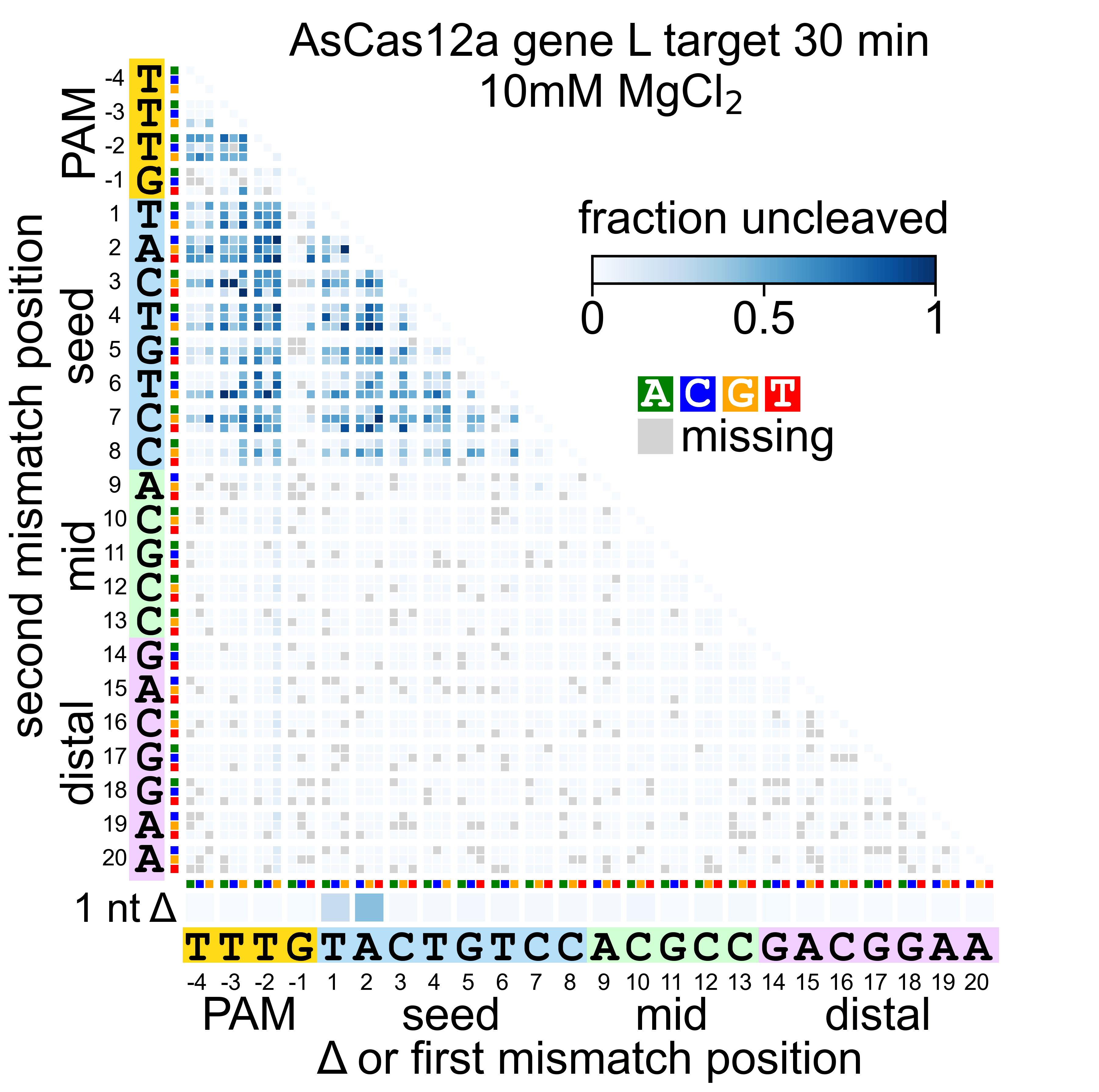

### As_W_1_nicked.gif

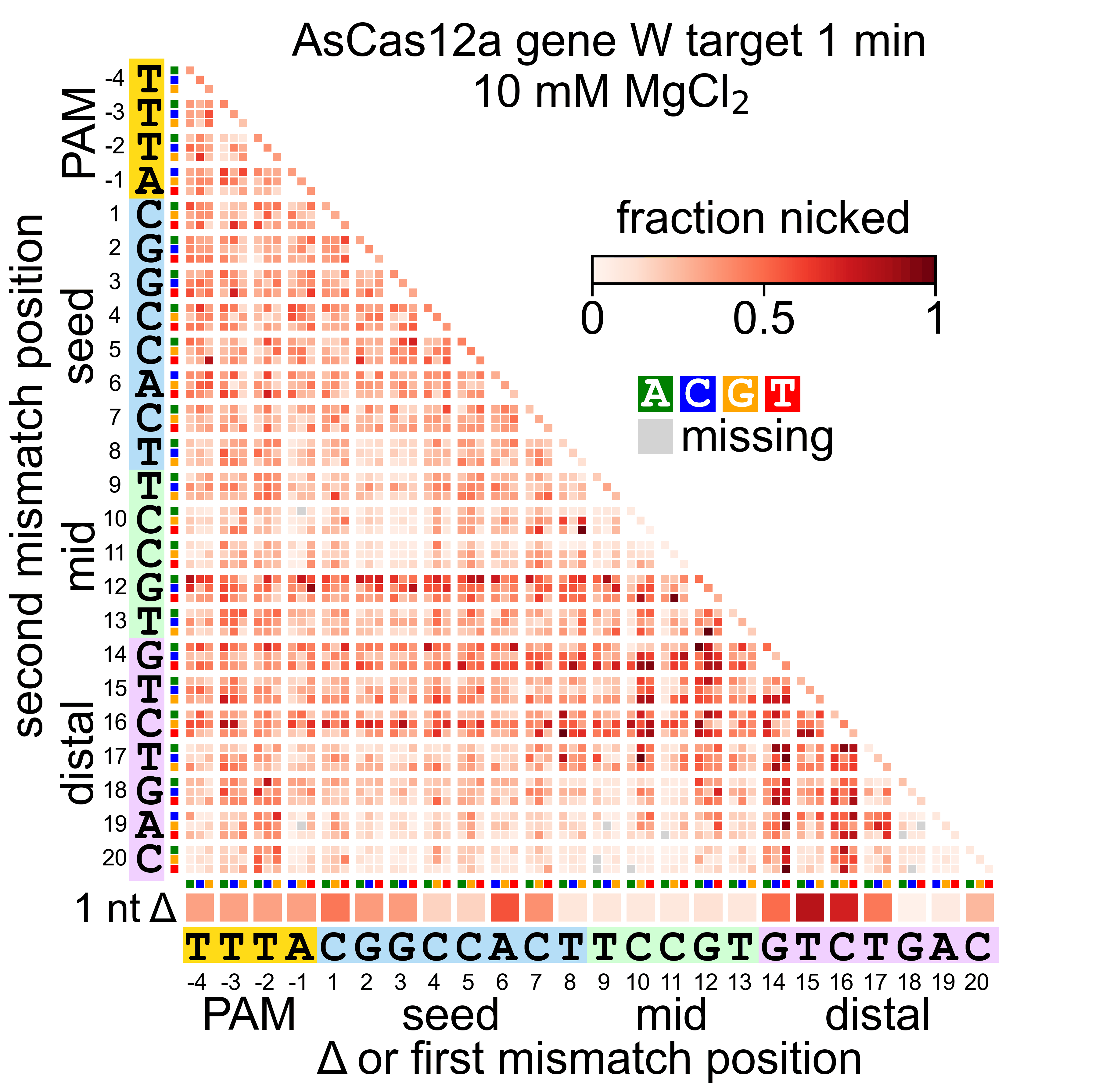

### As_W_1_uncleaved.gif

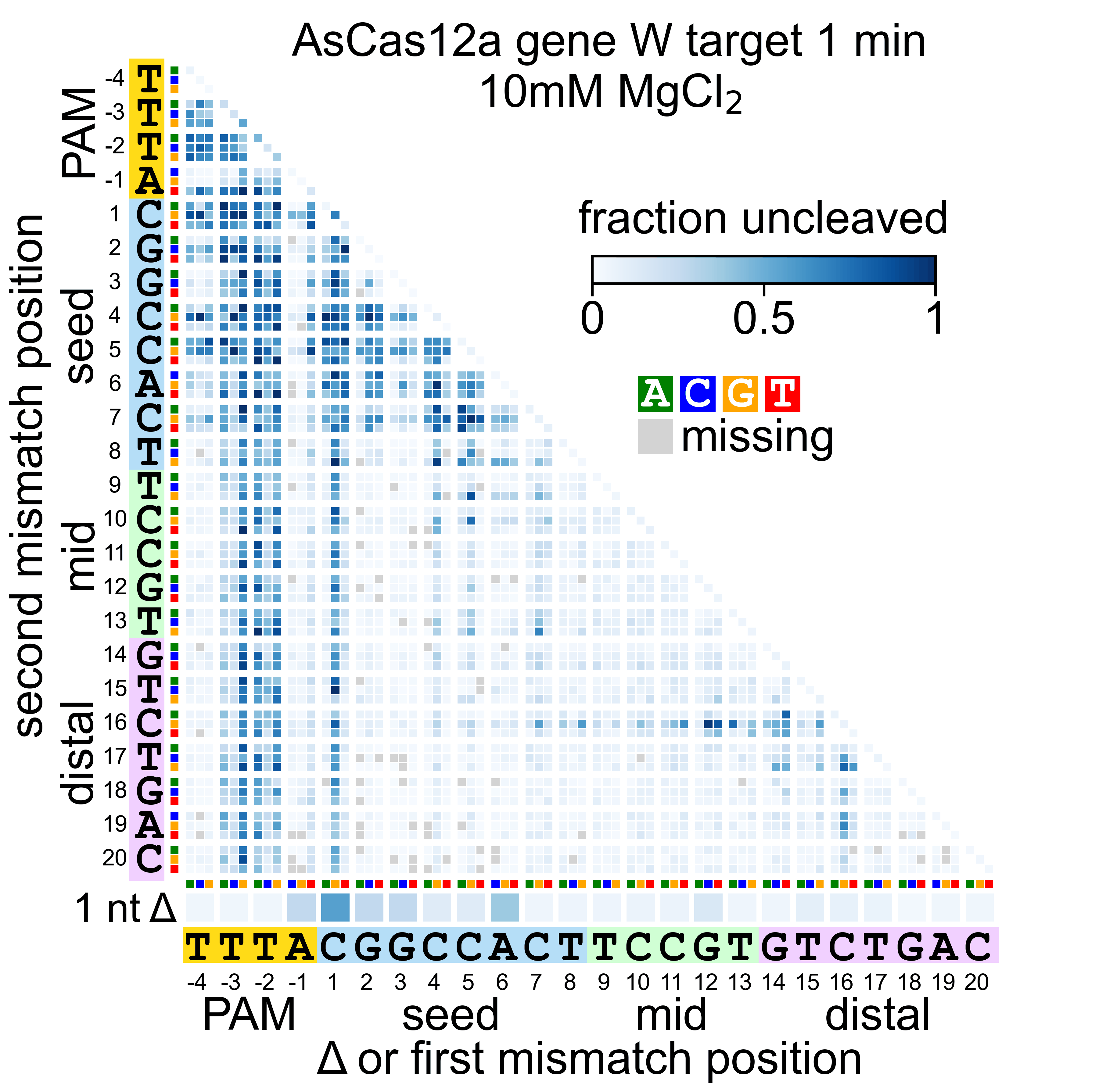

### As_W_30_nicked.gif

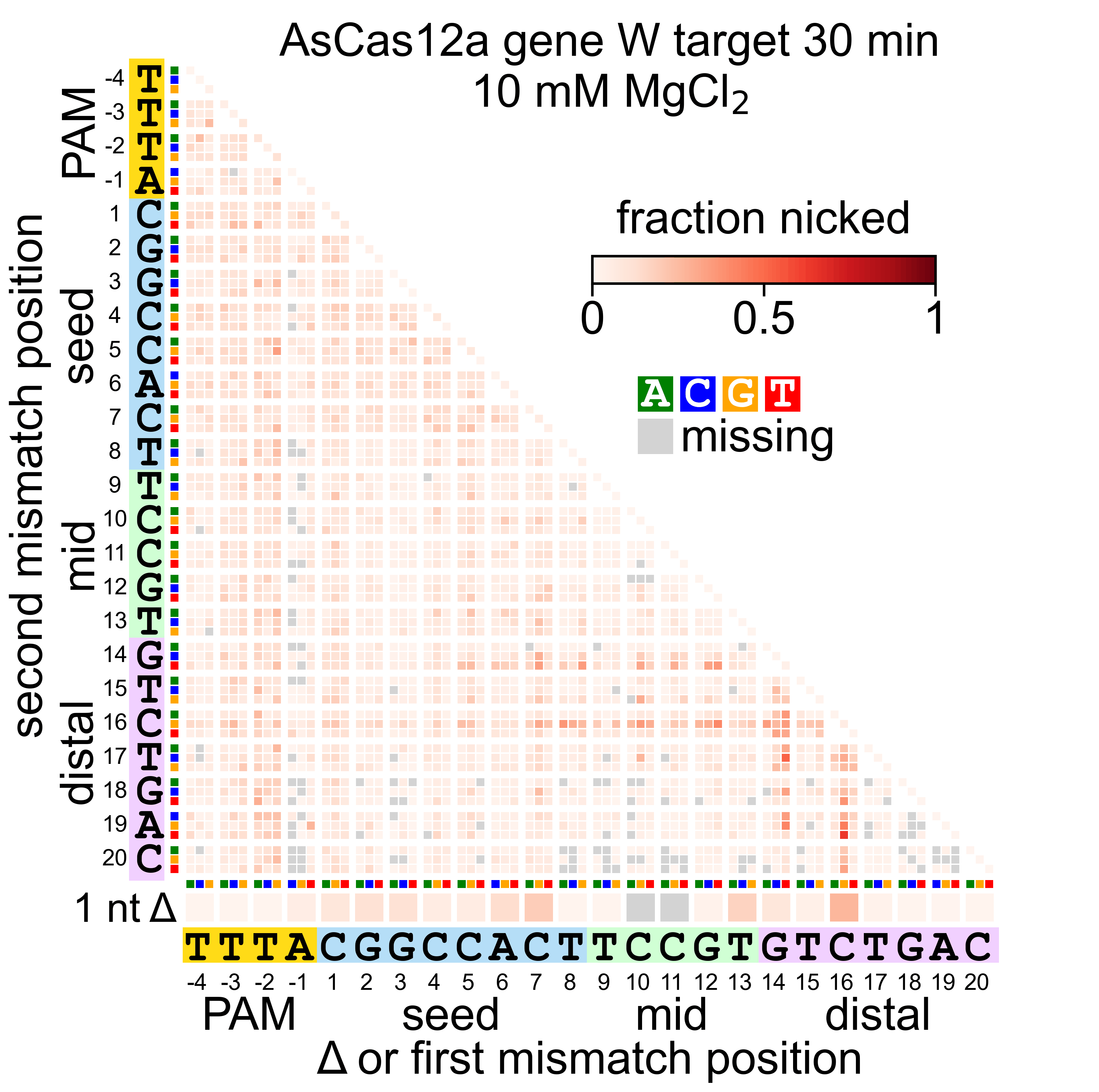

### As_W_30_uncleaved.gif

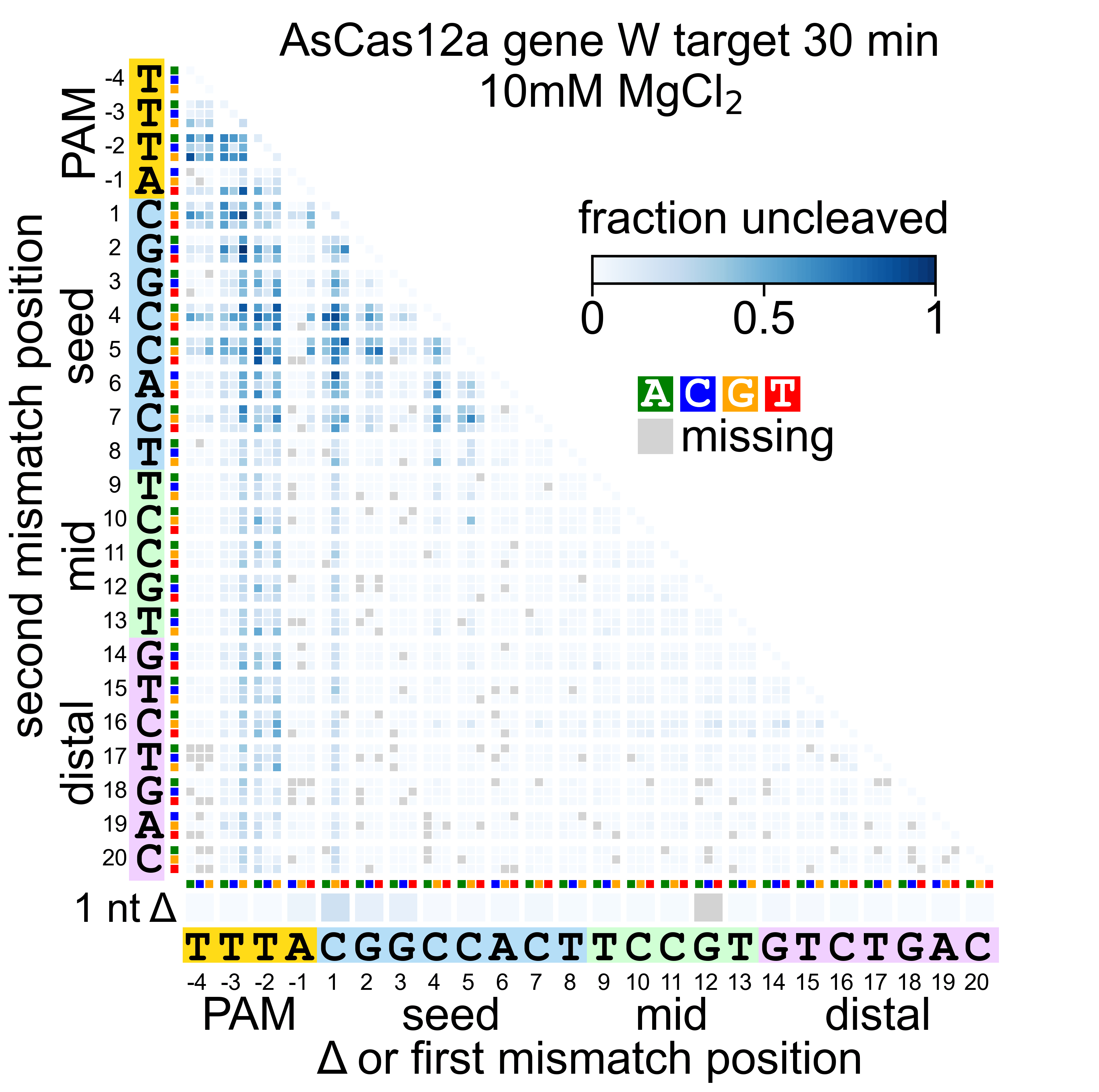

### Fn_L_1_nicked.gif

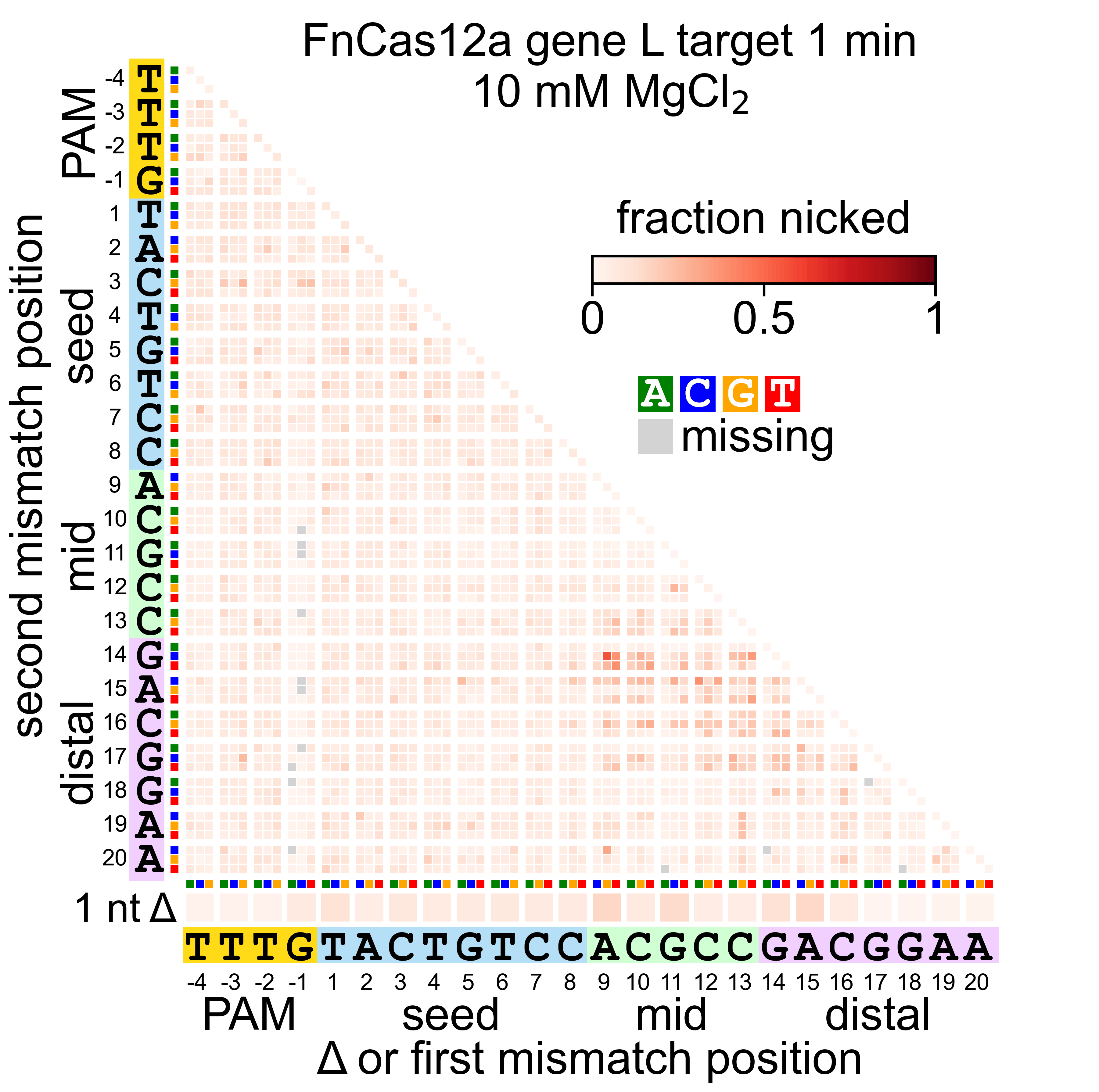

### Fn_L_1_uncleaved.gif

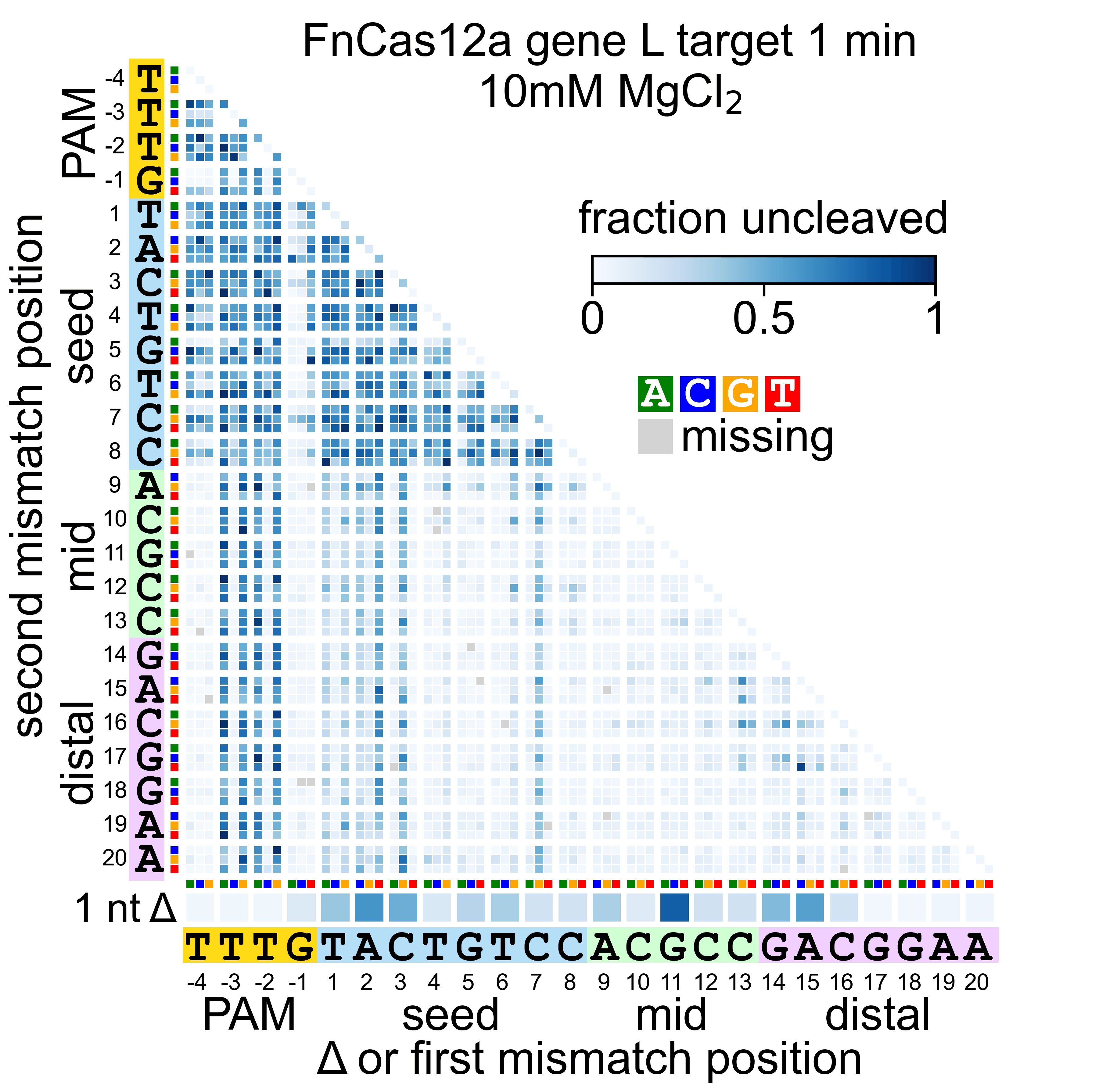

### Fn_L_30_nicked.gif

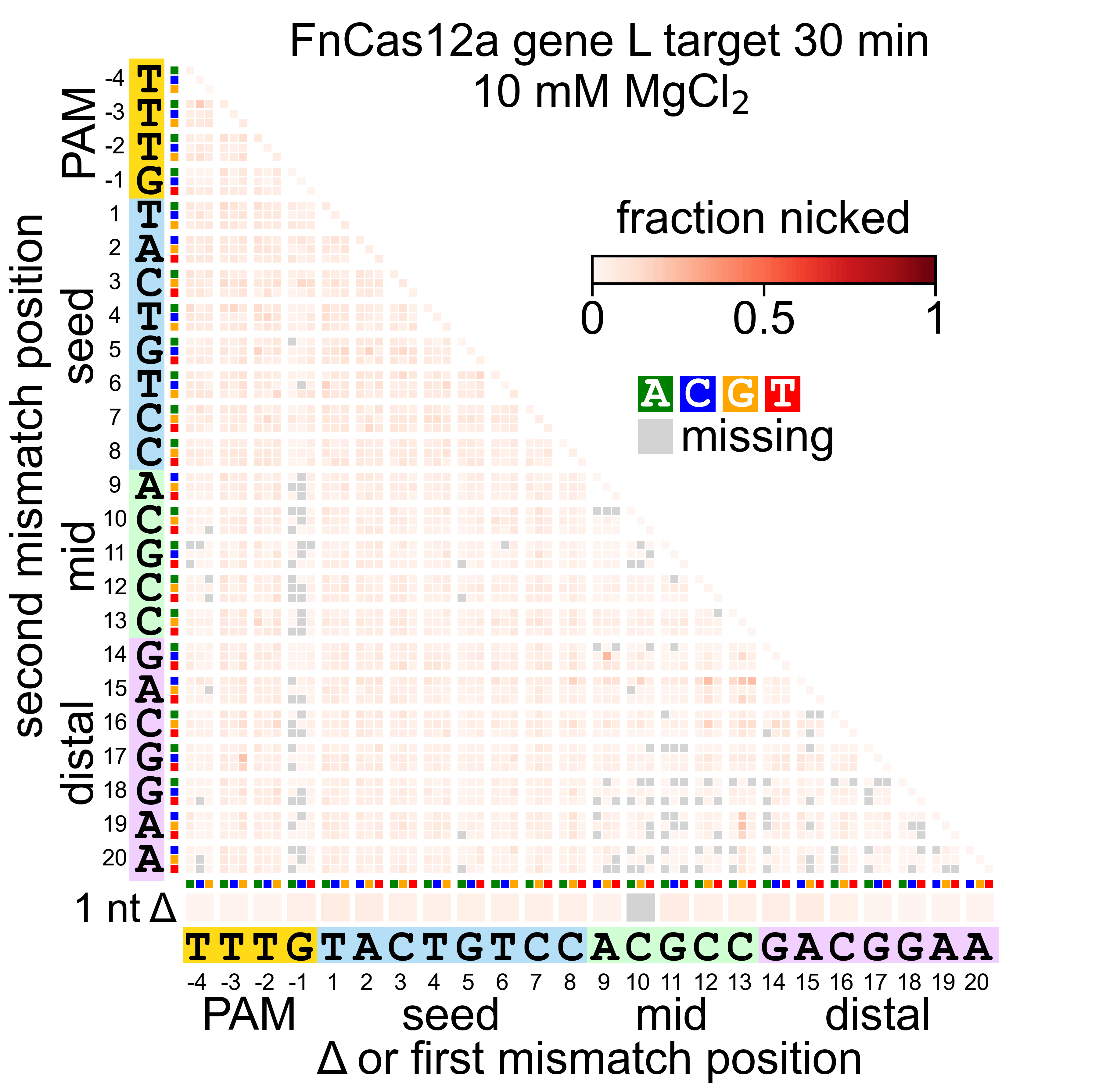

### Fn_L_30_uncleaved.gif

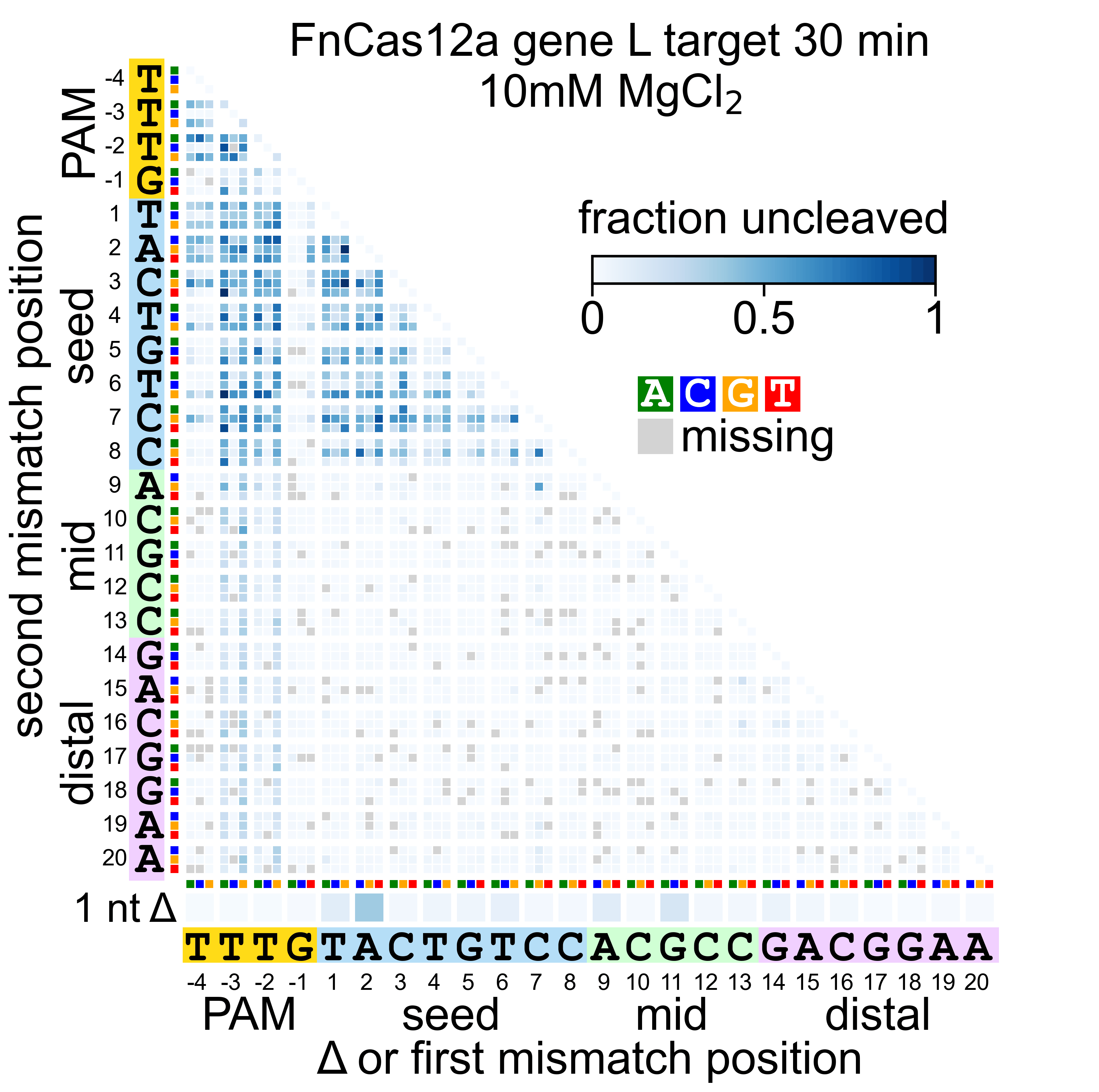

### Fn_W_1_nicked.gif

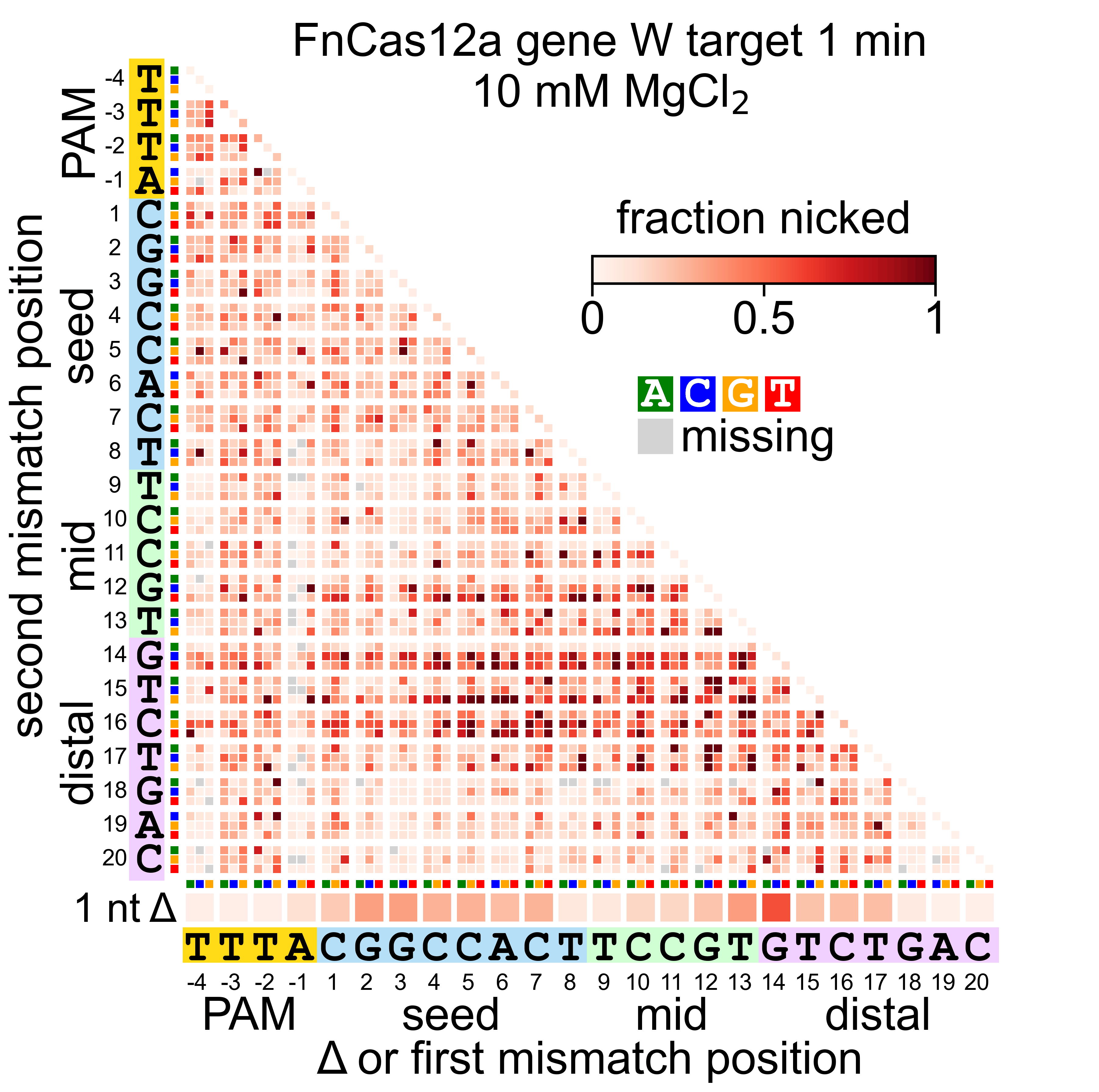

### Fn_W_1_uncleaved.gif

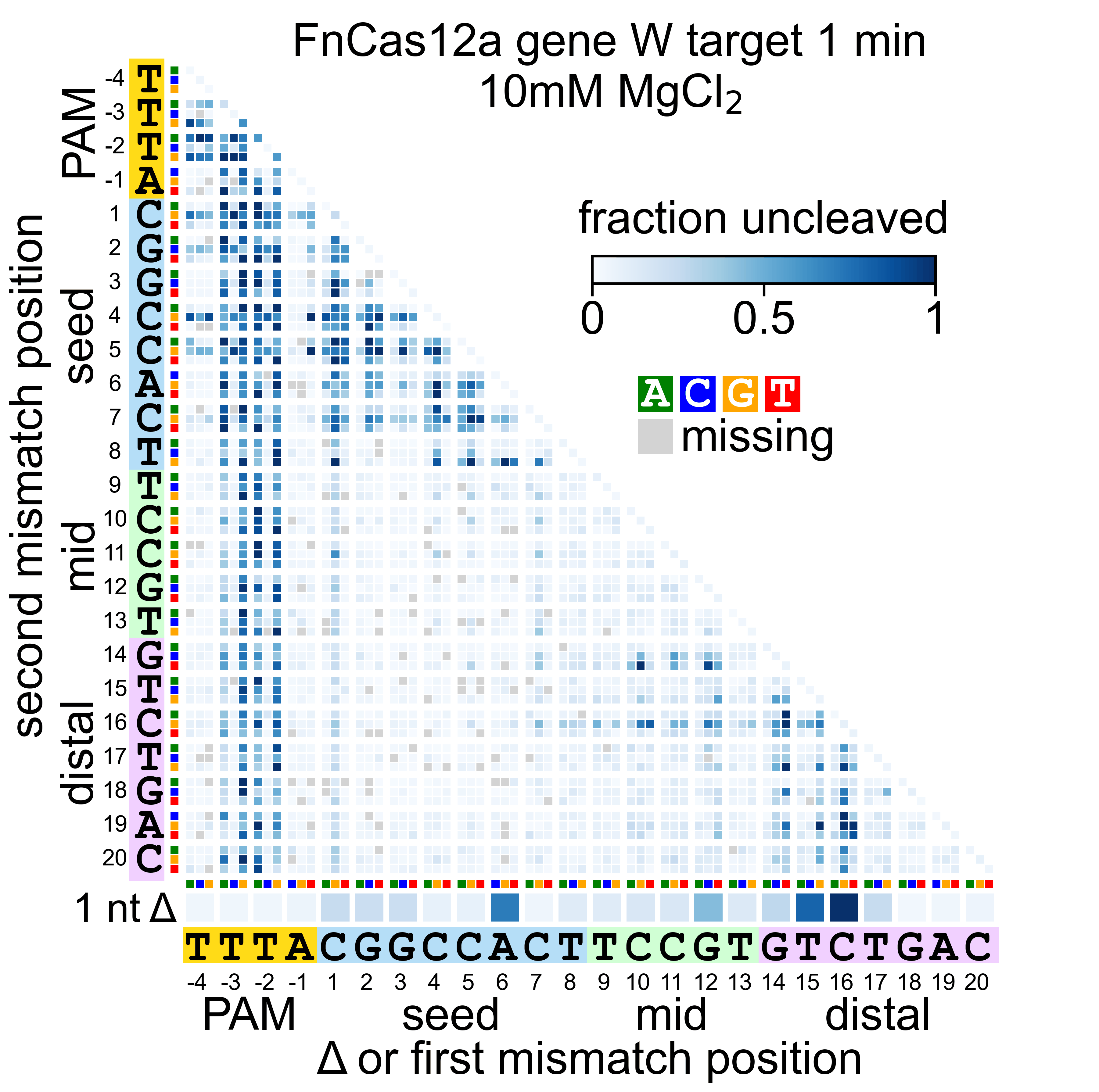

### Fn_W_30_nicked.gif

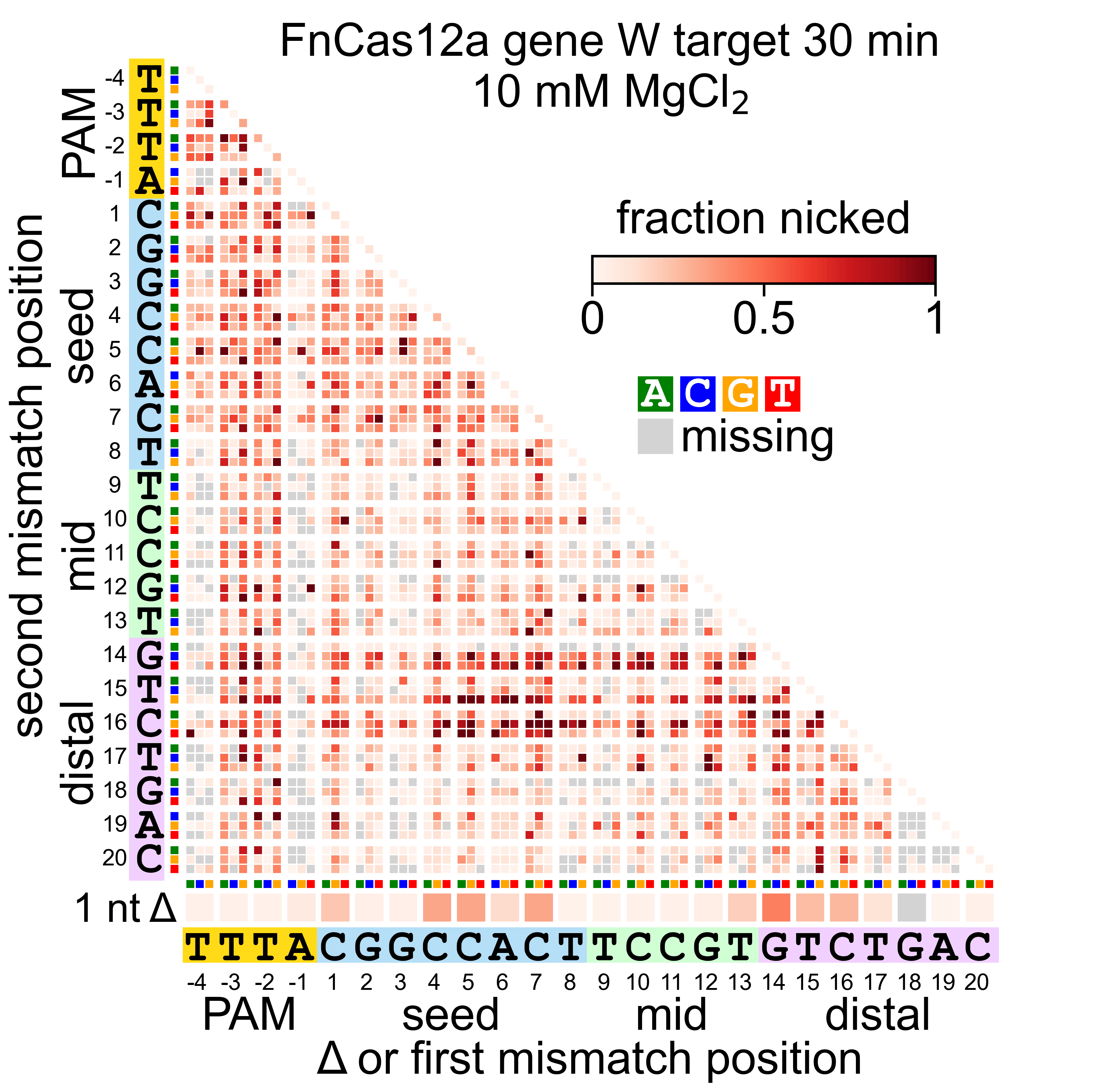

### Fn_W_30_uncleaved.gif

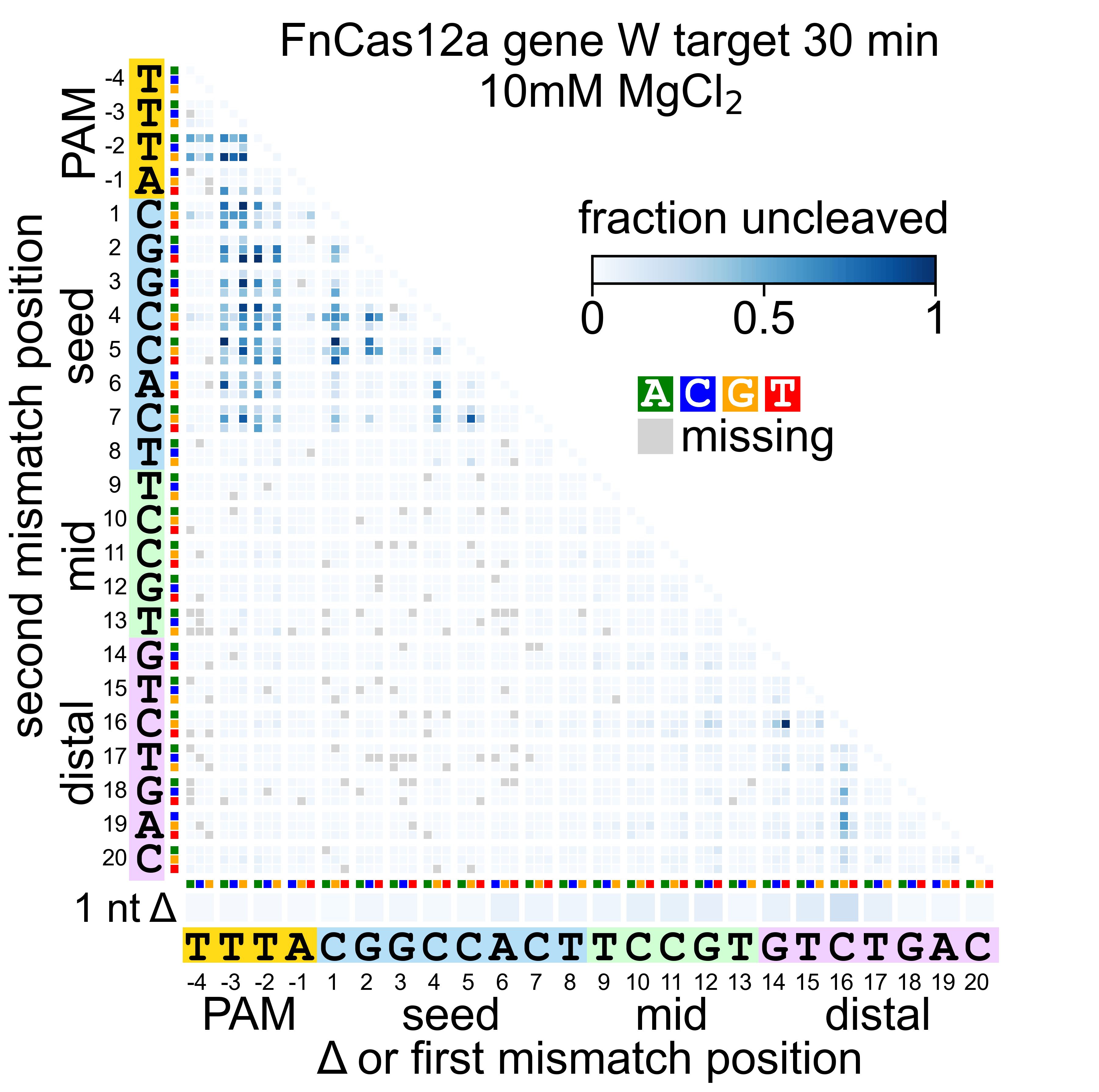

### Lb_L_1_nicked.gif

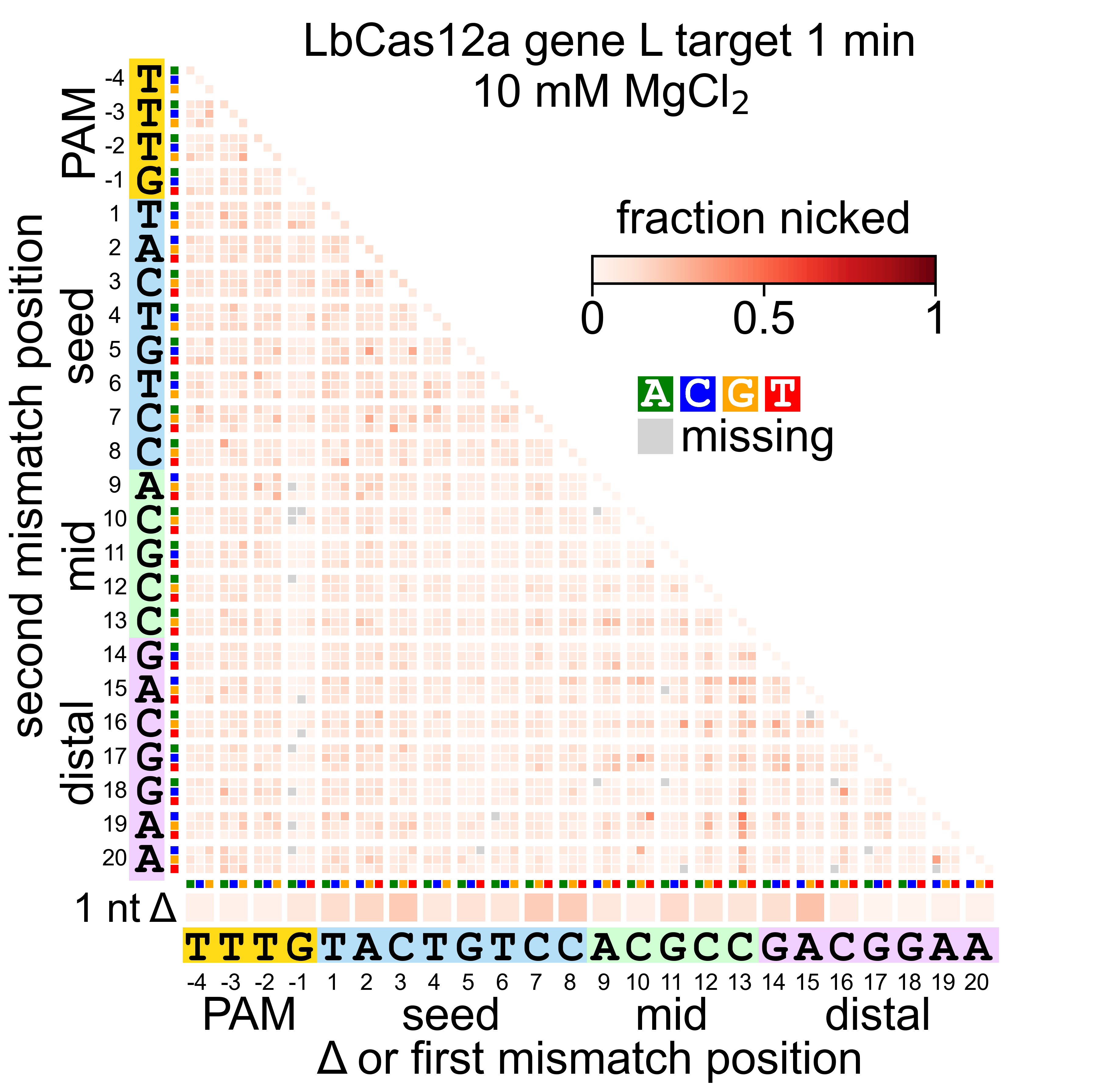

### Lb_L_1_uncleaved.gif

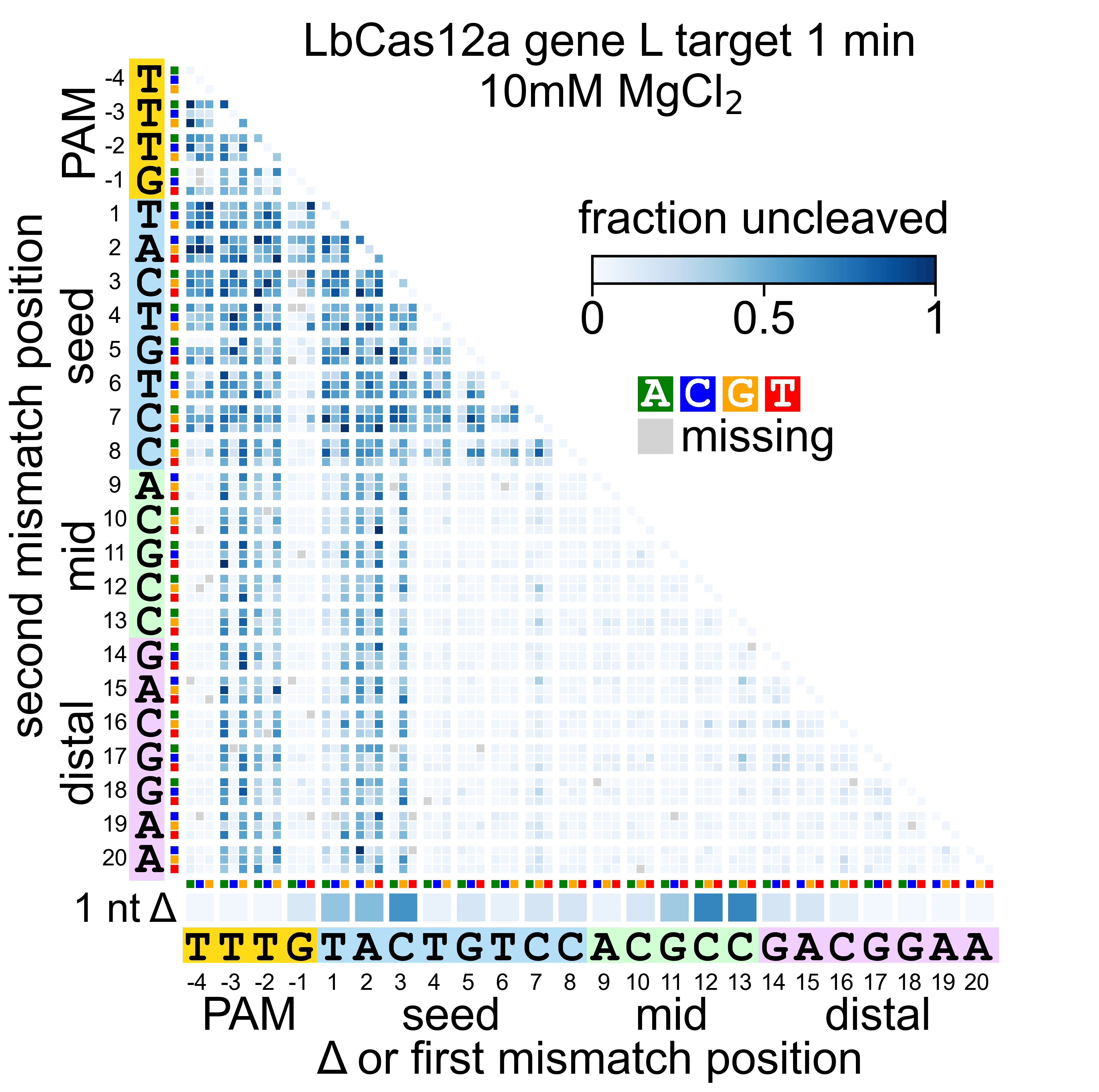

### Lb_L_30_nicked.gif

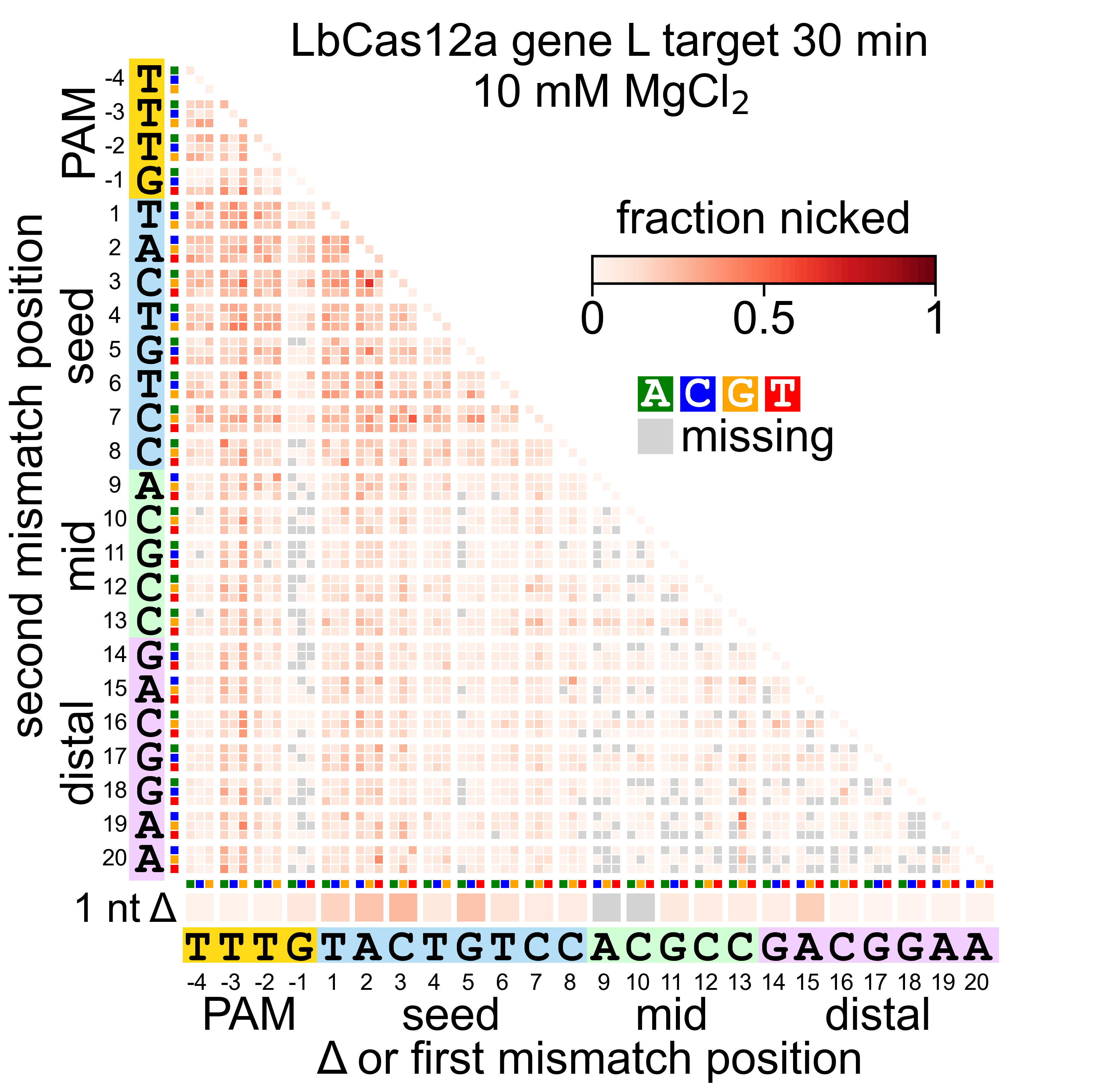

### Lb_L_30_uncleaved.gif

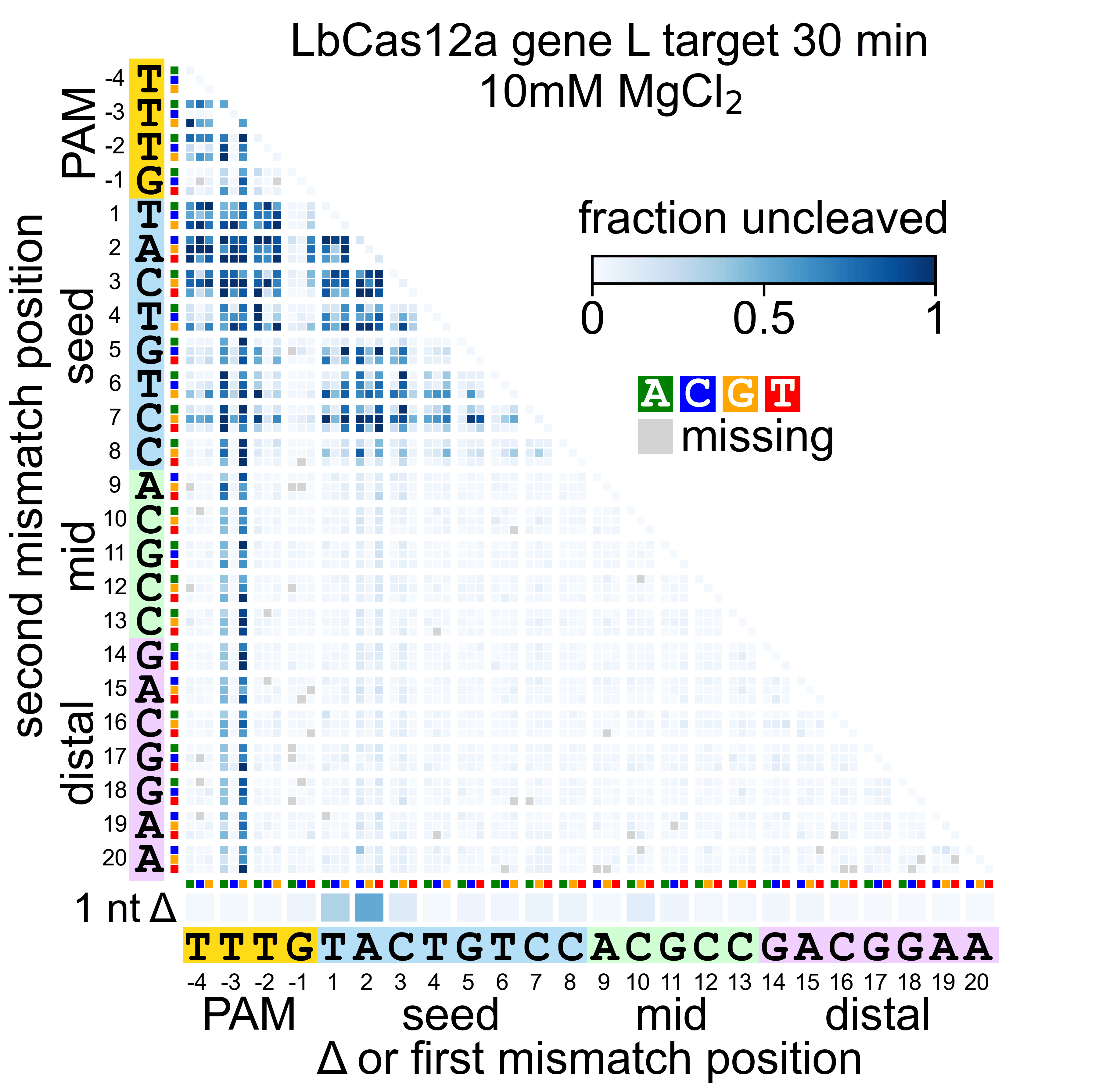

### Lb_W_1_nicked.gif

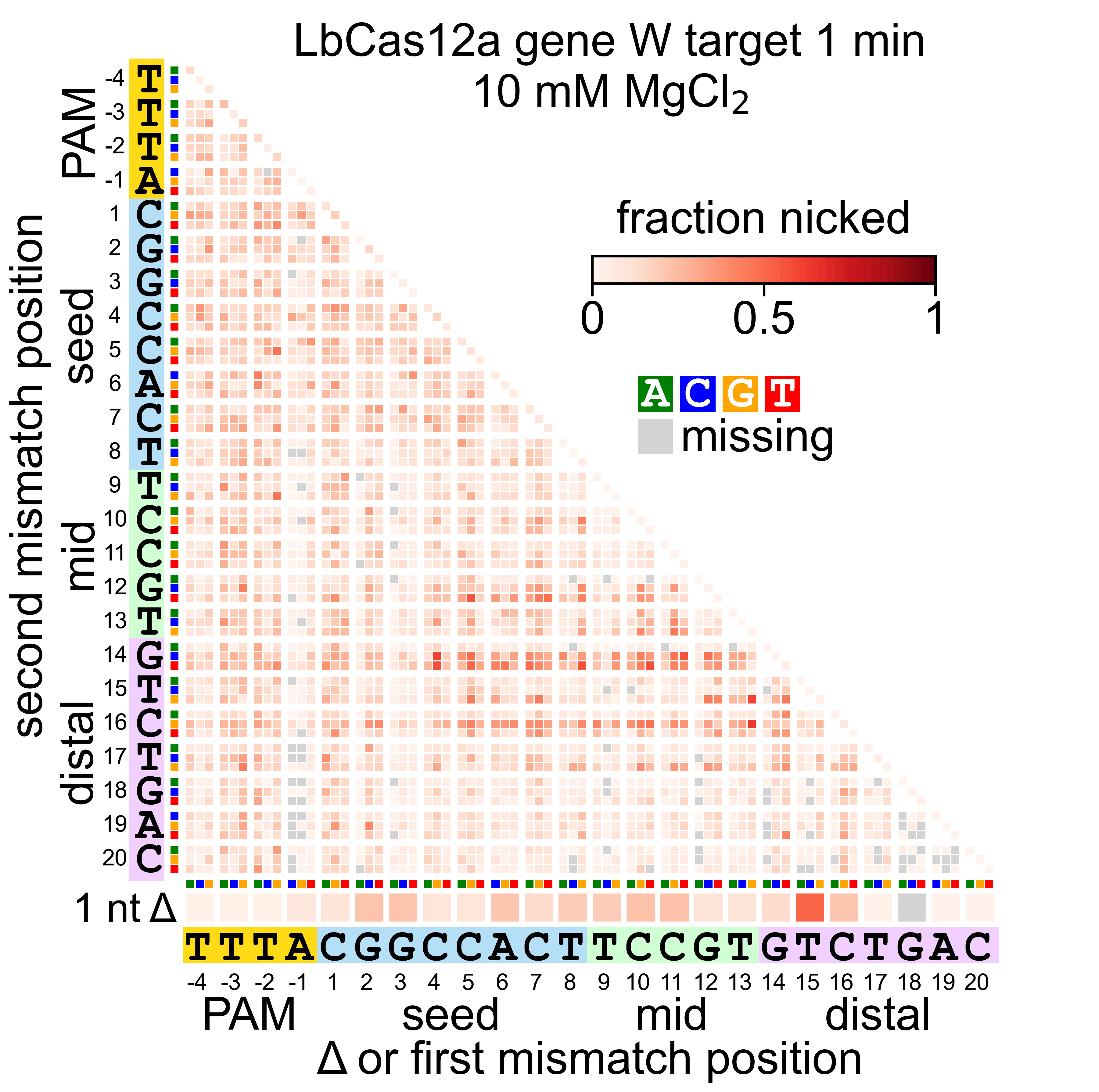

### Lb_W_1_uncleaved.gif

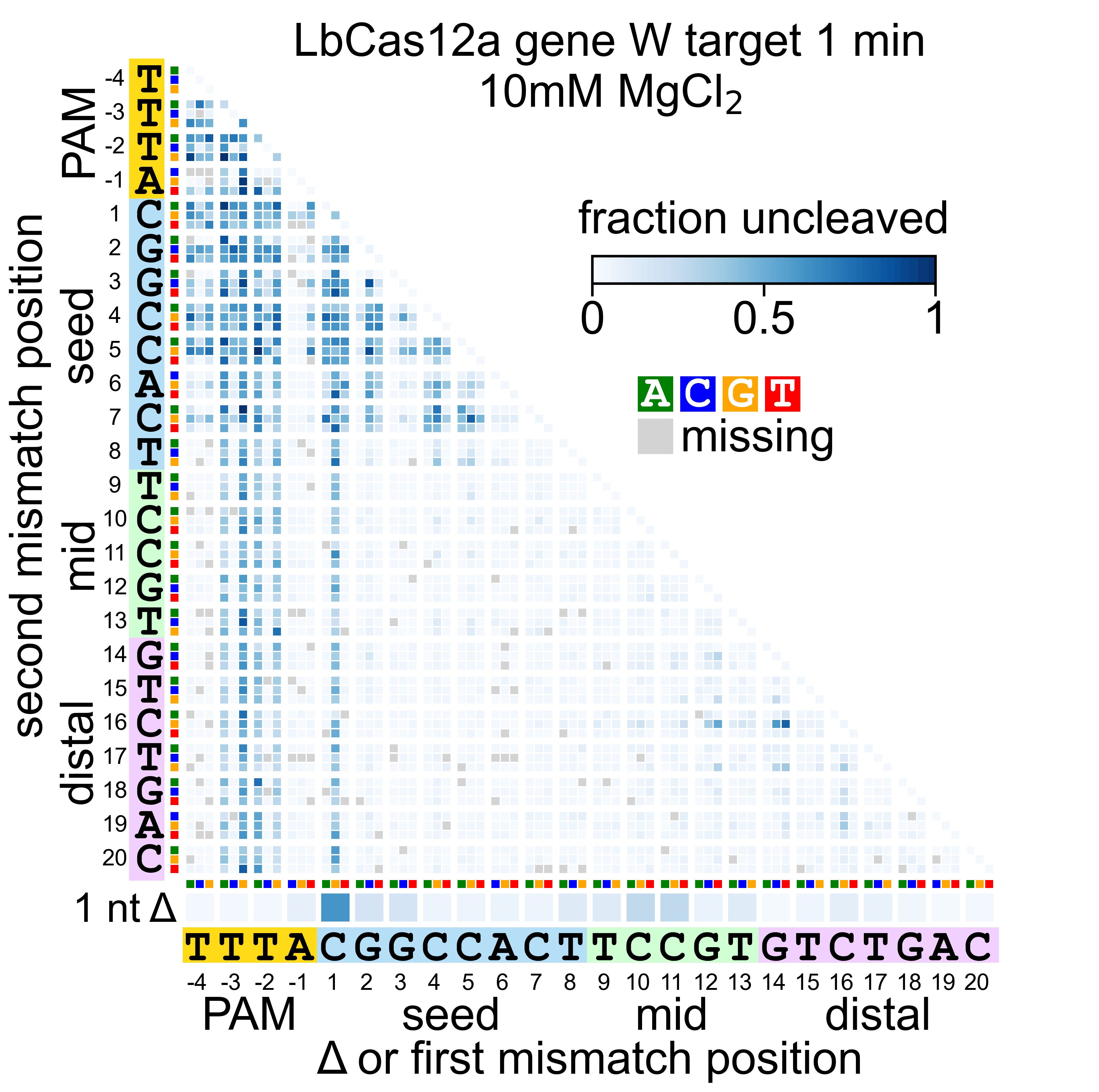

### Lb_W_30_nicked.gif

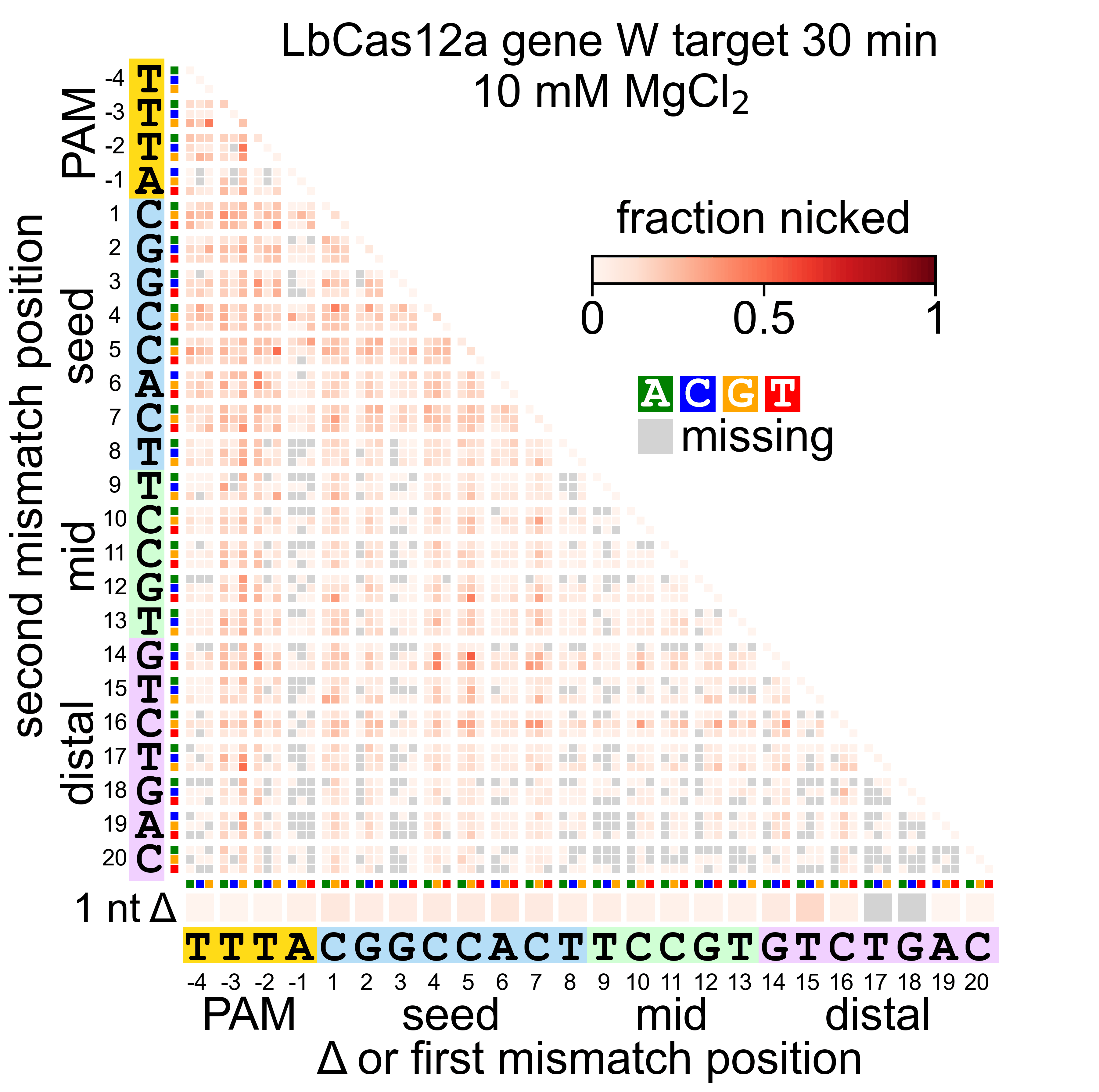

### Lb_W_30_uncleaved.gif

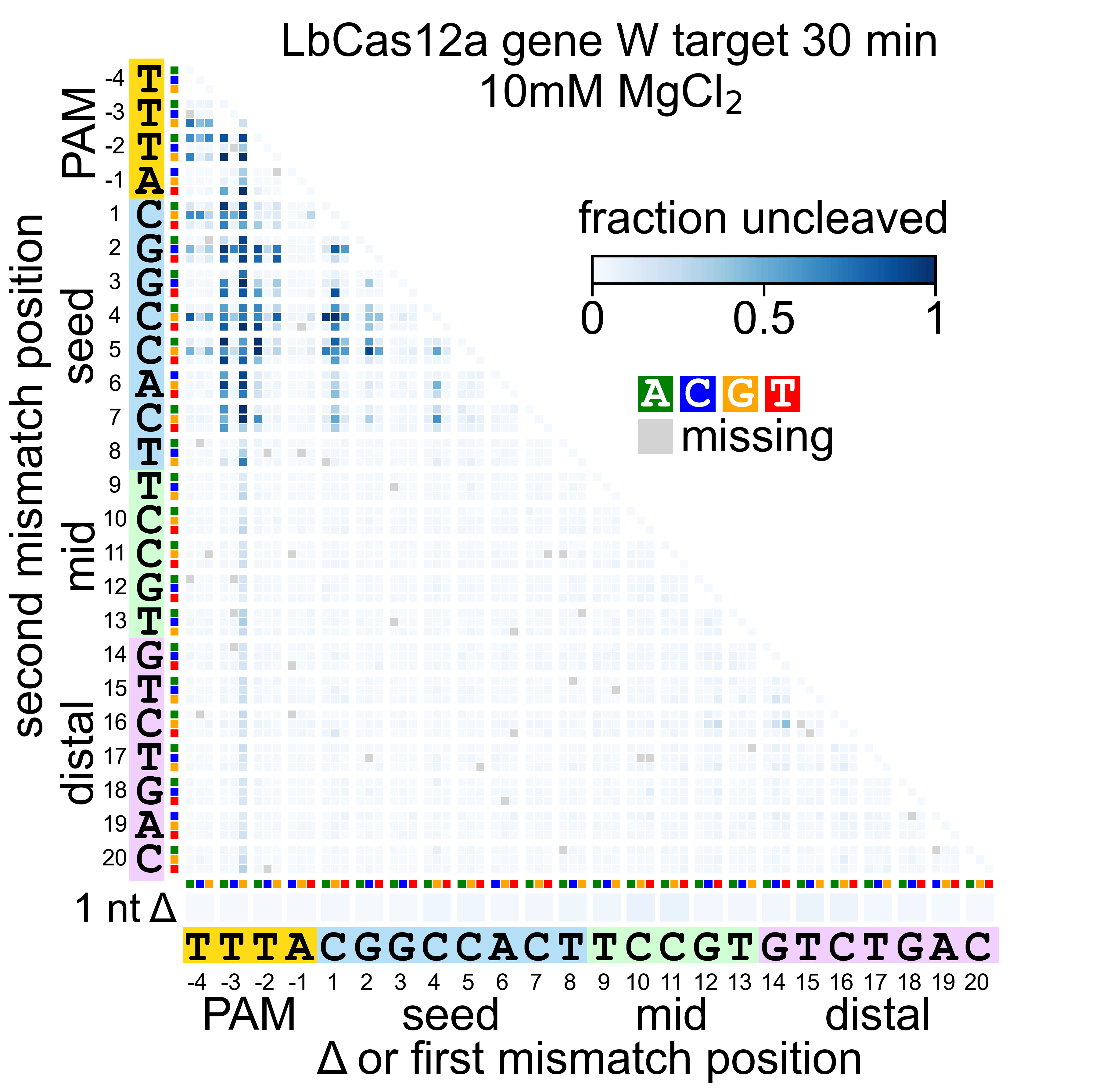
